# Bayesian mapping of the striatal microcircuit reveals robust asymmetries in the probabilities and distances of connections

**DOI:** 10.1101/2021.06.08.447507

**Authors:** François Cinotti, Mark D. Humphries

## Abstract

The striatum’s complex microcircuit is made by connections within and between its D1- and D2-receptor expressing projection neurons and at least five species of interneuron. Precise knowledge of this circuit is likely essential to understanding striatum’s functional roles and its dysfunction in a wide range of movement and cognitive disorders. We introduce here a Bayesian approach to mapping neuron connectivity using intracellular recording data, which lets us simultaneously evaluate the probability of connection between neuron types, the strength of evidence for it, and its dependence on distance. Using it to synthesise a complete map of the mouse striatum, we find strong evidence for two asymmetries: a selective asymmetry of projection neuron connections, with D2 neurons connecting twice as densely to other projection neurons than do D1 neurons, but neither subtype preferentially connecting to another; and a length-scale asymmetry, with interneuron connection probabilities remaining non-negligible at more than twice the distance of projection neuron connections. We further show our Bayesian approach can evaluate evidence for wiring changes, using data from the developing striatum and a mouse model of Huntington’s disease. By quantifying the uncertainty in our knowledge of the microcircuit, our approach reveals a wide range of potential striatal wiring diagrams consistent with current data.

## INTRODUCTION

As the input of the basal ganglia circuit, the striatum has been ascribed key computational roles in action selection (Redgrave et al., 1999; Gurney et al., 2001a; Liénard and Girard, 2014), decision making (Bogacz and Gurney, 2007; Ding and Gold, 2010, 2012; Yartsev et al., 2018), and reinforcement learning (Reynolds et al., 2001; Samejima et al., 2005; Bornstein and Daw, 2011; Khamassi and Humphries, 2012; Gurney et al., 2015). Within the striatum is a microcircuit comprising the GABAergic spiny projection neurons (SPNs), which make up to 97% of striatal neurons in the rat (Oorschot, 2013), and at least five species of predominantly GABAergic interneurons (Burke et al., 2017; Tepper et al., 2018). These SPNs divide into two populations that express either the D1 or D2-type of dopamine receptors (Gerfen et al., 1990; Gerfen and Surmeier, 2011). Projections from the D1 and D2 SPN populations respectively form the striatonigral and striatopallidal pathways (Gerfen et al., 1990; Kreitzer, 2009; Gerfen and Surmeier, 2011), through which they influence dynamics throughout the basal ganglia and beyond. The microcircuit’s connections onto the D1 and D2 SPNs are then a potentially major actor in sculpting the output of this nucleus, and thus the computations ascribed to it.

One key to understanding the role of the microcircuit in the computations of striatum is knowing the relative influence of one neuron type on another (Alexander and Wickens, 1993; Hjorth et al., 2009; Humphries et al., 2009; Lau et al., 2010; Ponzi and Wickens, 2010; Klaus et al., 2011; Damodaran et al., 2014). Two broad influences of this microcircuit on the output of SPNs are well-known: the feedforward inhibition by GABAergic interneurons, and feedback inhibition by lateral connections between the SPNs (Plenz, 2003; Tepper et al., 2004, 2008; Humphries et al., 2010). But to understand how all elements of the striatum’s microcircuit influence its output requires a full account of the microcircuit’s wiring, which we currently lack. To address this problem, here we synthesise data from pairwise intracellular recording studies to generate a statistically-rigorous and comprehensive map of the wiring probabilities between the key neuron species of the mouse striatum.

A key issue in estimating connection probabilities from intracellular recording data is that recording studies report a single probability for each connection type, given by the rate of successful connections between two types of neuron, without providing any measures of uncertainty. In this paper, we solve this problem by introducing a Bayesian approach to estimating the probability of connection between neuron types using pairwise intracellular recording data, which allows us to draw rigorous conclusions about the strength of evidence for claims about the microcircuit. Using this approach on data from the mouse striatum, we show that the previously reported asymmetry between the rates at which D1 and D2 neurons make connections is robust, with D2 SPNs having roughly twice the connection rate of D1 SPNs; but contrary to previous claims we also show there is no evidence for an asymmetry in the rates at which they receive connections, and so there is no preferential target for D1 or D2 SPNs. We then demonstrate a new method for using single measurements of connection rates to estimate distance-dependent probabilities and their uncertainty. Using these methods to analyse both SPN and interneuron connectivity, we complete our Bayesian map of the connectivity of the mouse striatum. Finally, we demonstrate how our Bayesian approach lets us quantify and test changes to that microcircuit map: we test the claim that D1 SPN connections are altered in a mouse model of Huntington’s disease, and find no evidence for it; and, using recent data from Krajeski et al. (2019), we show the selective asymmetry of D1 and D2 SPNs appears at different stages during development. Our Bayesian approach thus simultaneously evaluates the probability of connection between neuron types, its dependence on distance, and the strength of evidence for it, creating a solid foundation for theories of striatal computation.

## MATERIALS AND METHODS

### Data processing

We extracted data on pairwise connections from intracellular recordings of striatal neurons from a database of studies. The full set of data we extracted is given in Table 1. Because of the way Taverna et al. (2008) gave their results, namely reporting the number of connected pairs and specifying if any were bidirectional instead of reporting the number of connections, the number of tests for non-mixed pairs we use in this paper is doubled compared to the original study. For instance, when Taverna et al. (2008) say they found 5 connected pairs out of 19 pairs of D1 neurons, we interpret this as 5 connections out of 38 tests, to be consistent with the mixed D1 and D2 pairs which by necessity are unidirectional (a D1 → D2 connection can only be tested in one direction or it becomes a D2 → D1 connection). This was also the case for the data of Cepeda et al. (2013) on SPN connections in wild-type and Huntington’s disease model mice.

**Table 1.**
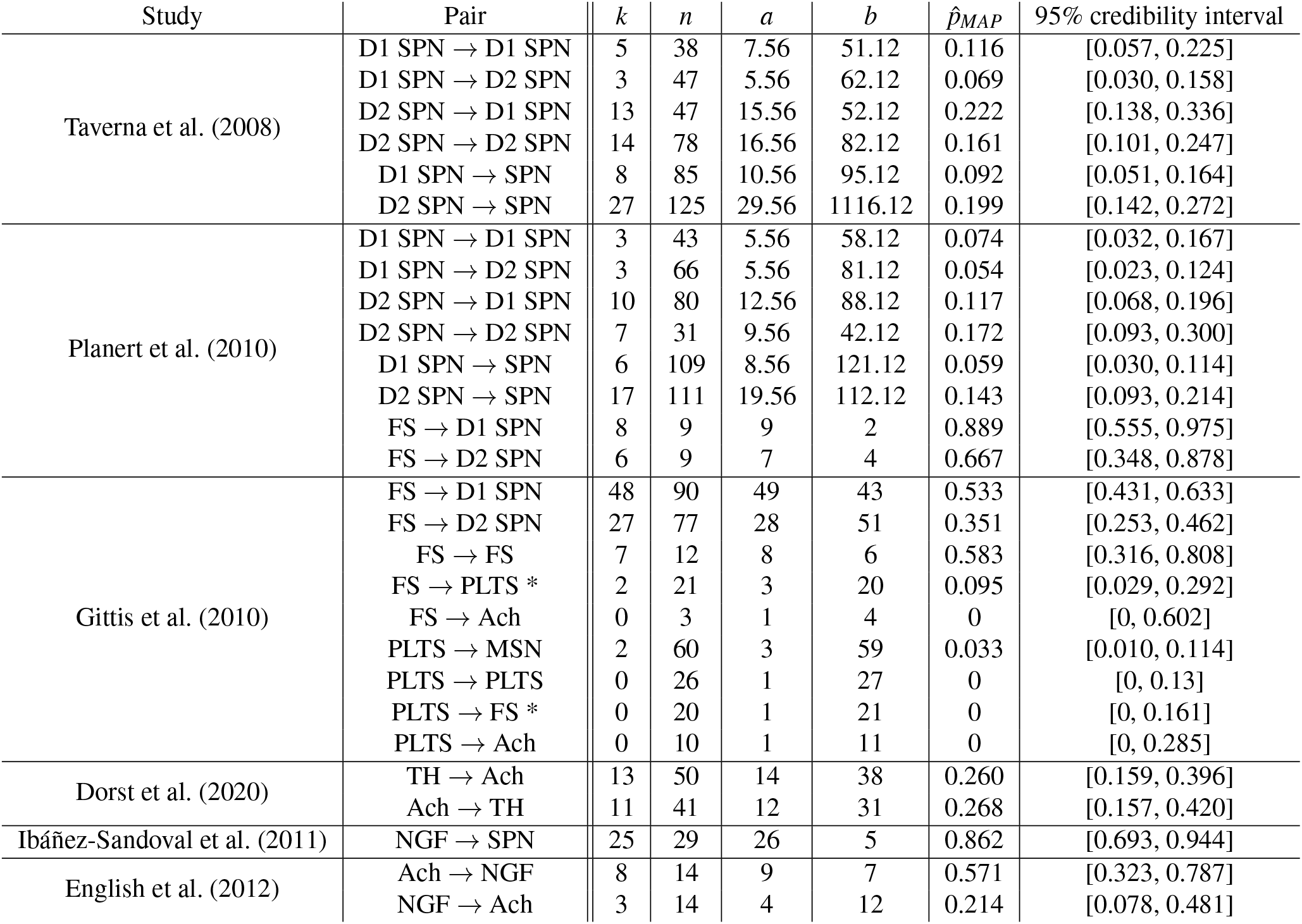
Pairwise connection data from mice used to build the Bayesian map of the striatum microcircuit, alongside Bayesian estimates of the connection probabilities. *k*: number of connected pairs in that study; *n* number of sampled pairs; *a, b* parameters of the resulting Beta distribution for the posterior of *p* using either the literature prior for SPN connections or a uniform prior for interneurons as explained in the main text; 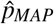, the MAP estimate of *p*. Abbreviations for neuron names: SPN: Spiny Projection Neuron; FS: Fast Spiking interneuron, PLTS: Persistent Low Threshold Spiking interneuron; Ach: Cholinergic interneuron; TH: Tyrosyne-Hydroxylase interneuron; NGF: (NPY-expressing) NeuroGliaForm interneuron. * Data concerning FS → PLTS and PLTS → FS connections from Gittis et al. (2010) was pooled with that of Szydlowski et al. (2013).

| Study | Pair | $k$ | $n$ | $a$ | $b$ | $\hat{p}_{MAP}$ | 95% credibility interval |
| --- | --- | --- | --- | --- | --- | --- | --- |
| Taverna et al. (2008) | D1 SPN → D1 SPN | 5 | 38 | 7.56 | 51.12 | 0.116 | [0.057, 0.225] |
| | D1 SPN → D2 SPN | 3 | 47 | 5.56 | 62.12 | 0.069 | [0.030, 0.158] |
| | D2 SPN → D1 SPN | 13 | 47 | 15.56 | 52.12 | 0.222 | [0.138, 0.336] |
| | D2 SPN → D2 SPN | 14 | 78 | 16.56 | 82.12 | 0.161 | [0.101, 0.247] |
| | D1 SPN → SPN | 8 | 85 | 10.56 | 95.12 | 0.092 | [0.051, 0.164] |
| | D2 SPN → SPN | 27 | 125 | 29.56 | 1116.12 | 0.199 | [0.142, 0.272] |
| Planert et al. (2010) | D1 SPN → D1 SPN | 3 | 43 | 5.56 | 58.12 | 0.074 | [0.032, 0.167] |
| | D1 SPN → D2 SPN | 3 | 66 | 5.56 | 81.12 | 0.054 | [0.023, 0.124] |
| | D2 SPN → D1 SPN | 10 | 80 | 12.56 | 88.12 | 0.117 | [0.068, 0.196] |
| | D2 SPN → D2 SPN | 7 | 31 | 9.56 | 42.12 | 0.172 | [0.093, 0.300] |
| | D1 SPN → SPN | 6 | 109 | 8.56 | 121.12 | 0.059 | [0.030, 0.114] |
| | D2 SPN → SPN | 17 | 111 | 19.56 | 112.12 | 0.143 | [0.093, 0.214] |
| | FS → D1 SPN | 8 | 9 | 9 | 2 | 0.889 | [0.555, 0.975] |
| | FS → D2 SPN | 6 | 9 | 7 | 4 | 0.667 | [0.348, 0.878] |
| Gittis et al. (2010) | FS → D1 SPN | 48 | 90 | 49 | 43 | 0.533 | [0.431, 0.633] |
| | FS → D2 SPN | 27 | 77 | 28 | 51 | 0.351 | [0.253, 0.462] |
| | FS → FS | 7 | 12 | 8 | 6 | 0.583 | [0.316, 0.808] |
| | FS → PLTS * | 2 | 21 | 3 | 20 | 0.095 | [0.029, 0.292] |
| | FS → Ach | 0 | 3 | 1 | 4 | 0 | [0, 0.602] |
| | PLTS → MSN | 2 | 60 | 3 | 59 | 0.033 | [0.010, 0.114] |
| | PLTS → PLTS | 0 | 26 | 1 | 27 | 0 | [0, 0.13] |
| | PLTS → FS * | 0 | 20 | 1 | 21 | 0 | [0, 0.161] |
| | PLTS → Ach | 0 | 10 | 1 | 11 | 0 | [0, 0.285] |
| Dorst et al. (2020) | TH → Ach | 13 | 50 | 14 | 38 | 0.260 | [0.159, 0.396] |
| | Ach → TH | 11 | 41 | 12 | 31 | 0.268 | [0.157, 0.420] |
| Ibáñez-Sandoval et al. (2011) | NGF → SPN | 25 | 29 | 26 | 5 | 0.862 | [0.693, 0.944] |
| English et al. (2012) | Ach → NGF | 8 | 14 | 9 | 7 | 0.571 | [0.323, 0.787] |
| | NGF → Ach | 3 | 14 | 4 | 12 | 0.214 | [0.078, 0.481] |

### Bayesian inference of connection probabilities

A single experimental test for determining whether one neuron connects to another will yield either a positive or negative result, so that it is equivalent to a Bernoulli test with a success rate *p*, the unknown probability of connection we are trying to infer. When analysing a whole study consisting of several of these tests, we assume that each test is independent and shares the same success rate *p* with the others. Thus, the study as a whole can be described using a binomial distribution:

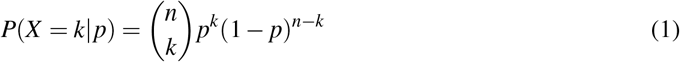

where *k* is the number of connected pairs and *n* the total number of tested pairs of that type. In this way, the binomial distribution provides a likelihood for the data given *p*.

Our goal is to estimate this *p*, the probability of connection, and the uncertainty of that estimate. According to Bayes theorem, the posterior distribution for *p* can be determined by:

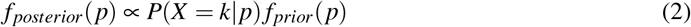

given a prior *f_prior_*(*p*), which is a probability distribution describing our initial beliefs about the possible value of *p*. Finding a posterior for the success rate of a binomial distribution is a well known problem in Bayesian inference and the prior distribution used in this case is a beta distribution:

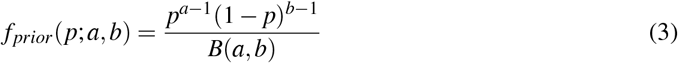

with *a* and *b* the parameters determining the shape of the prior, and *B*(*a, b*) the so-called Beta function. The main advantage of this type of prior, known as the conjugate prior of binomial distributions, is that the posterior that results from combining this prior with a likelihood in the form of a binomial distribution simply turns out to be a new beta distribution with updated parameters (sparing us the trouble of renormalising the right hand side of equation 2 to get a proper probability density function):

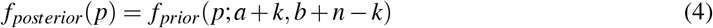

In other words, to determine the posterior we simply have to add the number of successful tests *k* to *a*, and the number of unsuccessful tests *n* – *k* to *b*.

Consequently, obtaining the posterior distribution is a single line of code. In MATLAB, this is `posterior = betapdf(p, a + k, b + n-k)` with *p* a vector of probabilities of connection for which we want the corresponding probability density value, and *a* and *b* the shape parameters of the initial prior.

### Design of the prior based on previous literature

In the Results, we test a set of standard priors for the Beta distribution, the uniform prior (*a* = *b* = 1), the Jeffreys prior (*a* = *b* = 0.5), and the Haldane prior (*a* = *b* = 0). We also test a prior based on previous literature of connections between SPNs, which we derive here. Knowing its mean *μ* and variance *v*, the shape parameters of a beta distribution are:

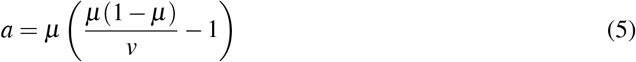

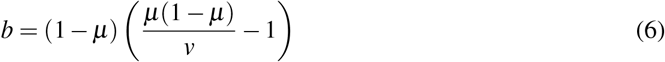

Previous studies that did not differentiate the D1 and D2 subtypes have shown that SPNs connect to one another at a mean rate of 0.12 (Czubayko and Plenz, 2002; Tunstall et al., 2002; Koos et al., 2004; Taverna et al., 2004), leaving us with a decision to make about the desired variance of the prior. Despite their thoroughness (325 tested pairs in Koos et al. (2004)), we could not directly use a beta distribution based on the number of pairs in the initial studies, as the resulting variance, which would reflect uncertainty attached to the measurement of the average connection rate between all types of pairs, would be so small that the new evidence with SPN subtype distinction would be unable to significantly affect the posterior. Indeed, the desired variance should reflect the fact that the average connection rate of 0.12 masks the potential existence of four distinct connection rates for each pair. We were unable to find a principled way of deriving this desired variance and, for this reason, different values of variance were tested before settling for 0.005 which gives the corresponding beta distribution a shape that makes such a prior both sufficiently informative as to be interesting without being completely insensitive to the addition of new data. Setting *μ* = 0.12 and *v* = 0.005, we find *a* = 2.56 and *b* = 18.12.

### Inferring distance-dependent probabilities of connection from point estimates

Intracellular recording studies typically report a maximum distance of pairwise recording, so our point estimate *p* of the probability of connection is then actually an integral over any distance-dependent probability of connection. We show here how we can derive estimates for the distance-dependent probability of connection from these point estimates, using simple models.

We assume that the probability of connection from a source neuron to a target neuron at distance *r* away is an exponentially decreasing function of distance:

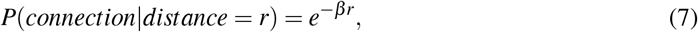

with decay parameter *β*. While a simple model, its advantage for us is its dependence on a single parameter *β*, which we show below can be inferred directly from our point estimate *p*, giving us a full posterior distribution for *β* too. Thus, while the model for *P*(*connection|distance*) is user-defined, we use our Bayesian inference approach to both fit the model’s parameter and obtain its uncertainty (indeed our approach is sufficiently general that any one parameter model could be used for *P*(*connection|distance*)).

Our goal here is to estimate the length scale of the decay of connectivity, particularly so that we may compare the scales between different types of connection, rather than find a detailed model of the distance-dependence decay of the probability of connections. Finding the most accurate models would require both having the exact distances between all pairs of sampled neurons (for example, all pairs of D1 SPNs sampled), which are often not readily available, and solving a range of issues, including: finding suitable models to fit the data; finding appropriate methods to fit the models to the data; determining whether the data has sufficient power to fit each model; determining whether the data has sufficient power to decide between different models; and determining whether the data has sufficient power to accurately recover the parameters of each model. The specific distance-dependent model of a particular type of connection in the striatum is thus a considerable piece of work, beyond our scope here. Moreover, it is unlikely in any case to markedly change the estimated length-scale over which the probability of connection decays, as we expect distance-dependence to decay exponentially: models of connectivity in the striatum derived from overlapping models of dendrites and axons (Humphries et al., 2010; Hjorth et al., 2020) and data from cortical slices (Levy and Reyes, 2012) and cultures (Barral and Reyes, 2016) all show that probabilities of connection between neurons exponentially decrease with distance. Our exponential model is thus a reasonable choice.

Given our model for *P*(*connection|distance*), we now want to find the mapping *p*(*β*) between a given point-estimate probability of connection *p* and the decay parameter *β*. The mapping between *p* and *β* can be expressed as:

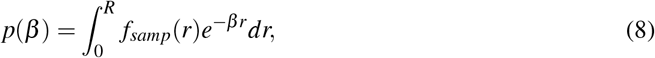

which is the product of the probability *f_samp_*(*r*) of experimenters selecting a neuron at distance *r* from another, and of the probability of these neurons being connected knowing *r* (equation 7), integrated over all possible values of *r* (see Figure 3 C for a visual depiction of what equation 8 means). *R* is the maximum distance at which the experimenters recorded their pairs of neurons.

Taking a central neuron as a reference point, we start by looking for a distribution for *r*, the distance between that central neuron and other neurons chosen for testing. By definition of a probability density function, *f_samp_*(*r*) must be such that the probability that *r* is found between two arbitrary values *r*_1_ and *r*_1_ + Δ*r* is:

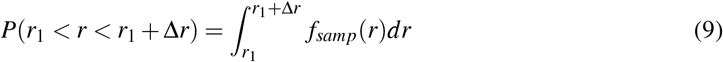

This probability distribution for distance depends on how the experimenters sampled their pairs. We consider two models for *f_samp_*(*r*). Our first model is that, given a starting neuron, experimenters are equally likely to sample any target neuron within their maximum recording radius – we call this model *f_equi_*. Our second model is that, from the starting neuron, experimenters will sample its nearest neighbour – we call this model *f_NN_*. Next we derive the *f_equi_* model, and describe the *f_NN_* model below.

### The equiprobable sampling model

For a given distance *r*_1_ from a central neuron, the probability of selecting a neuron within the volume bounded by *r*_1_ and *r*_1_ + Δ*r* is also equal to the ratio of the expected number of neurons found within it over the expected number of neurons in the total volume:

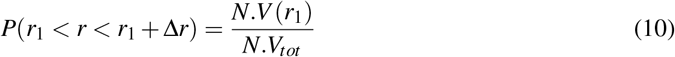

with *V*(*r*_1_) the subvolume bounded by *r*_1_ and *r*_1_ + Δ*r*, *V_tot_* the total volume and *N* the density of SPNs of whichever given type experimenters are currently trying to sample. Note that *N* cancels out in the fraction, which implies that the probability distribution for distance is, counter-intuitively perhaps, independent of post-synaptic SPN subtype, as long as the density is constant everywhere. According to the reported methods of the two studies we evaluate for SPN connections, experimenters selected neurons within the same field of focus at a maximum distance *R* of either 50*μm* (Taverna et al., 2008) or 100*μm* (Planert et al., 2010) which means that the total volume of interest is a cylinder of height *h*, corresponding to the depth of the field of focus, and the subvolume is a hollow cylinder, as depicted in Figure 3B:

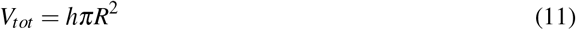

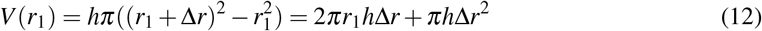

If we now combine the general definition of a probability density function (equation 9) with this particular equiprobable sampling assumption, we now have:

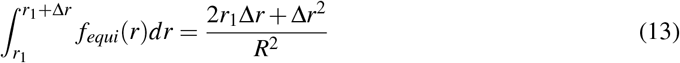

which we can solve to find *f_equi_* by differentiating the right hand side of the equation to obtain:

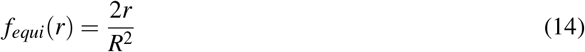

So that finally, by plugging equation 14 into equation 8 we obtain:

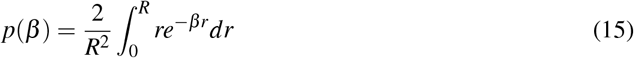

which can be used to create a mapping from *β* to *p, p*(*β*).

### The nearest-neighbour sampling model

To derive the nearest-neighbour model, we will consider the case where experimenters only patched pairs of neurons, ignoring the fact that Planert et al. (2010) would also patch triplets or quadruplets, and we will assume that they always patched the closest neuron within the maximum distance they set themselves. This means that we are looking for the density function for the nearest neighbour, *f_NN_*(*r*).

Because information about the nearest neighbour distribution was hard to find, we reproduce its derivation here, basing ourselves on Krider and Kehoe (2004), in case such a derivation would be of interest to others. Such a density function must satisfy:

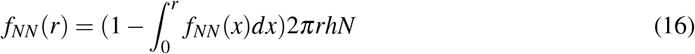

which states that the probability density that the nearest neuron is found at distance *r* is the product of the probability that the first neuron is not in fact found at a shorter distance from the central neuron (the first element in brackets on the right hand side) and of the probability that there is a neuron between *r* and *r* + *dr*. This latter probability is itself the product of the infinitesimal cylindrical volume found between *r* and *r* + *dr*, i.e. 2*πrhdr* with *h* (*μm*) the height of the cylinder, and *N* (*μm*^-3^) the average density of neurons. If we now differentiate *f_NN_*(*r*), we get:

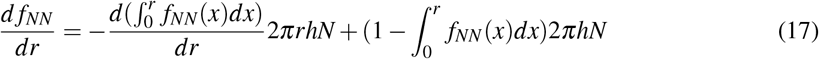

According to Leibniz’ rule, we get on the one hand:

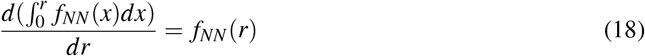

We can also substitute the term in brackets in the second half of equation 17, thanks to the following rewriting of equation 16:

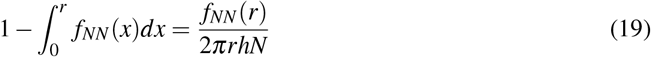

After these substitutions and factoring by *f_NN_*, we can now rewrite equation 17:

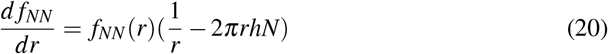

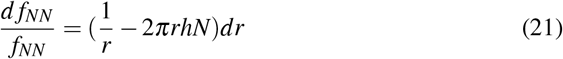

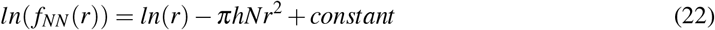

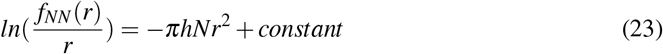

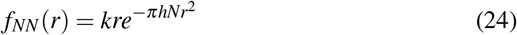

with *k*, the normalisation constant. Usually, *k* is defined so that integrating *f_NN_* between 0 and infinity is equal to 1, i.e. the nearest neighbour must be somewhere in that interval. In our particular case however, experimenters set themselves a maximum distance of either 50 or 100*μm*, meaning that the closest neuron must be closer than this distance (if there was no neuron closer than this, experimenters would simply look for another pair). In other words, *k* is such that:

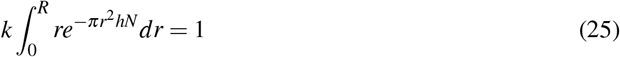

which ultimately gives us:

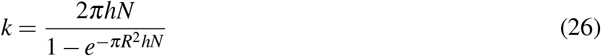

As previously, we can now combine these probabilities of sampling a neuron at a given distance with the probability of connection given distance (Equation 8) to find the overall probability of connection:

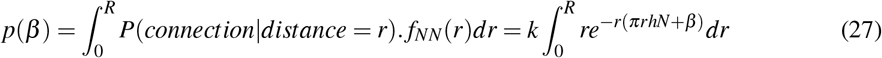

which gives us a new mapping between *p* and *β*.

Unlike the equiprobable model, the nearest-neighbours model depends on both the density of neurons *N* and the depth of the sampling plane *h*. Given that we have collapsed probabilities of connection based on the nature of the presynaptic neuron, we simply use an overall SPN density in the mouse brain of 80500 per mm^3^ following the convention of Hjorth et al. (2020) who chose this number based on the work of Rosen and Williams (2001) (and which is close to the estimated density of 84900 per mm^3^ in the rat brain (Oorschot, 1996)). As for *h*, the experimenters tell us that neurons were sampled in the same field of focus which would correspond to a height with an order of magnitude of a tenth or even a hundredth of micrometer. However, given that for a neuron to be in the same field of focus as another, it suffices that some part of its soma, whose diameter is between 10 and 20 *μm* in mice according to Gagnon et al. (2017), intersects this very small volume, we can expect *h* to be much larger in practice. Because of this uncertainty, we used three different values of *h* to get three different nearest-neighbour distributions: 0.1 *μm*, 1 *μm* and 10 *μm*.

### Transformation of posterior distributions

Having obtained the mapping *p*(*β*) between *p* and *β*, we can go a step further and find a full distribution (a posterior) for *β*, by transforming the posteriors we have previously obtained for *p*, *f_p_*(*p*), into posteriors for *β*, *f_β_*(*β*). By definition of a density function, for any possible values *a* and *b* of *β*, we have:

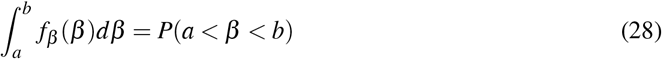

Thanks to the mapping from *β* to *p* (which is monotonically decreasing), we can also write:

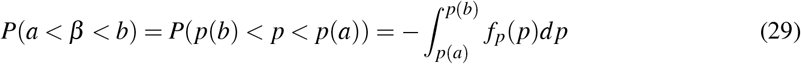

Finally, integration by substitution tells us:

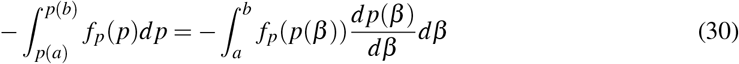

Hence, by identification with equation 28:

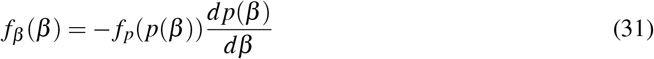

In order to draw *f_β_*, we converted regularly interpolated values of *β* into the corresponding values of *p* using equation 15 for the equiprobable sampling model or equation 27 for the nearest-neighbour model.

Obtaining the derivative of *p* with respect to *β* in equation 31 depended on the sampling process. In the case of equiprobable sampling, after an integration by parts of equation 15, we arrive at the following expression of *p*(*β*):

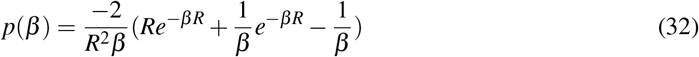

which can be differentiated with respect to *β*:

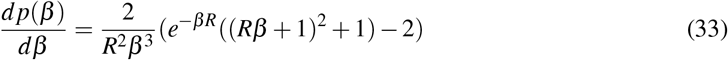

However, in the case of the nearest-neighbour distribution, we were unable to find a closed-form for 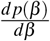, and resorted to a numerical approximation based on the regularly interpolated values of *β* and the corresponding values of *p* given by equation 27.

### Code availability

All code used for this work was written with MATLAB. The code necessary for the Bayesian analysis of *p*, the transformation of *f_p_*(*p*) to *f_β_*(*β*) and the Monte Carlo simulations depicted in Figure 4 C and D is available on the Github account of the Humphries lab: https://github.com/Humphries-Lab/Bayesian-map-of-striatum-circuitry.

## RESULTS

### Patch clamp data on connection rates between SPNs

We begin by reviewing key data on the connections within and between the D1- and D2-type SPNs, which we will also use to motivate our Bayesian approach. Previous studies by Taverna et al. (2008) and Planert et al. (2010) collected data on pairwise connections between SPN subtypes in slices obtained from both the dorsal and ventral striatum of mice. The subtype of the SPNs was determined by targeted expression of EGFP under the control of either a D1 or D2 receptor promoter sequence for Taverna et al. (2008) and of only a D1 receptor promoter for Planert et al. (2010). Non-labelled SPNs were then assumed to belong to whichever group was not meant to be labelled in this particular animal and electrophysiological criteria were used to exclude interneurons. After choosing a pair of neighbouring cells, no further apart than 50 *μm*, Taverna et al. (2008) detected connections in current-clamp mode by injecting a depolarising current step in the first – potentially presynaptic – neuron of the pair, and measuring depolarising postsynaptic potentials in the second one. This procedure was then repeated the other way around. Planert et al. (2010) recorded up to four neighbouring neurons simultaneously, with a larger maximum intersomatic distance of 100 *μm*, and detected connections by stimulating one neuron with a train of depolarising pulses to generate action potentials in the presynaptic neuron and recording the potential postsynaptic responses of the other neurons.

In both studies, the ratio of successful tests to the total number of tests (Table 1) was then reported as an estimate 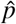 for the true probability of connection between these types of neuron (plotted as the bars in Figure 1A and B), in accordance with frequentist inference about a proportion. As they are estimates, they come with a level of uncertainty about the true proportion that depends on sample size, which here is the number of pairs that were tested. In a frequentist approach, this uncertainty would usually be given by a confidence interval.

**Figure 1.**
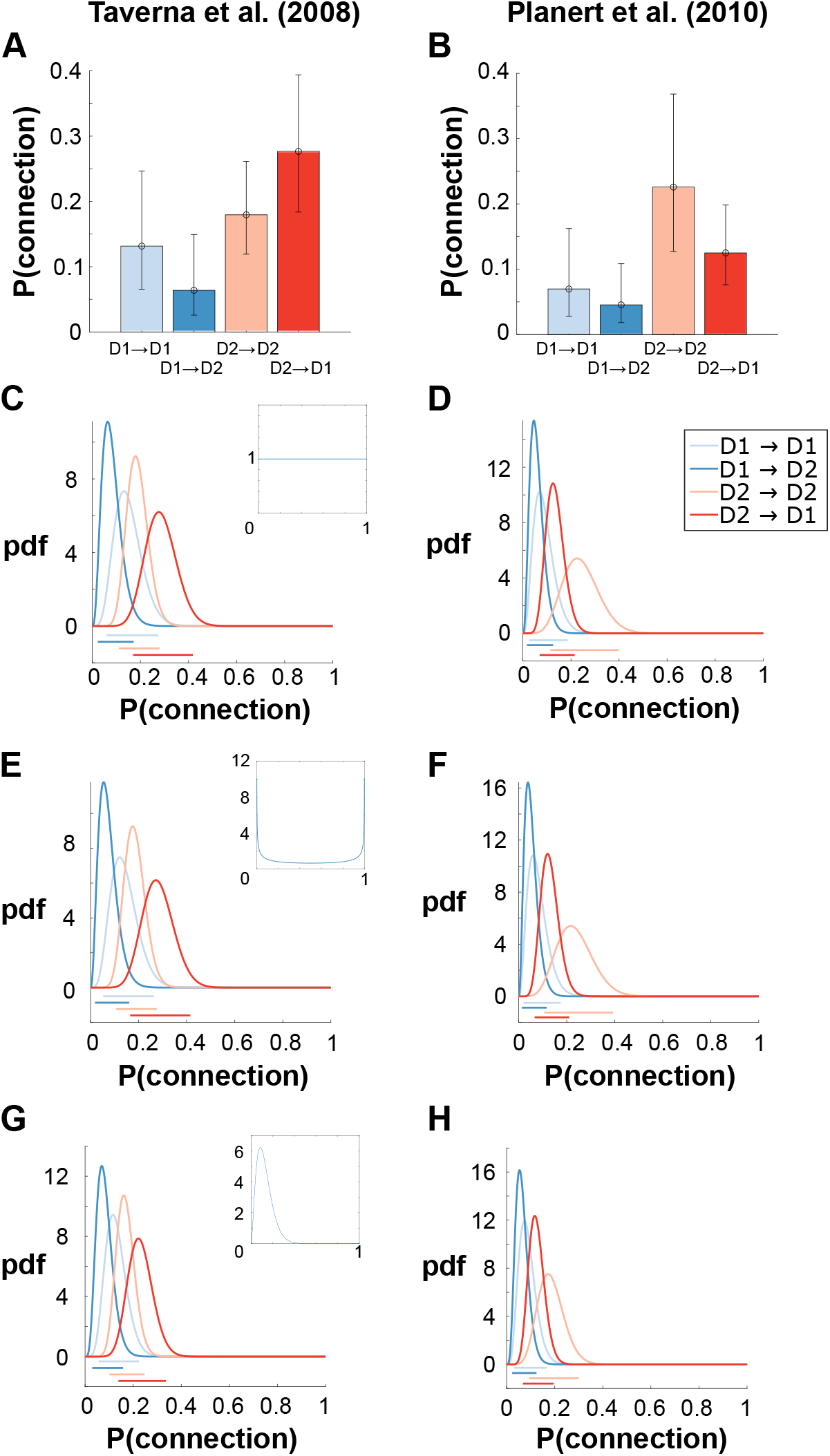
Probability of lateral connections between SPNs estimated using either frequentist or Bayesian methods. **A-B** Frequentist estimates of the probabilities of connection computed from intracellular recording data, and our computed 95% Wilson confidence intervals. **C-D** Posterior probability density functions for the probability of connection using a Bayesian approach. Coloured bars underneath the plot represent the 95% credibility intervals corresponding to each probability density function. Inset: shape of the prior, a uniform distribution. **E-F** Posterior probability density functions using the Jeffreys prior. **G-H** Posterior probability density functions using a prior based on previous literature with mean equal to 0.12 and variance equal to 0.005.

However, typically, intracellular recording studies do not report any estimate for the uncertainty surrounding their measurements of 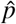, and recent theoretical studies of striatum (Burke et al., 2017; Hjorth et al., 2020; Bahuguna et al., 2015) have simply used the raw values 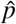 from Taverna et al. (2008) and Planert et al. (2010) to construct their models, supposing in particular that the probability of D1 to D1 connections is about twice as likely as D1 to D2 connections. When we do compute confidence intervals, such as the Wilson confidence interval for binomial proportions (Brown et al., 2001) that we add ourselves in Figure 1A and B, we find that, given the relatively small sample sizes, the confidence intervals overlap considerably.

### Bayesian inference of connection probabilities

As we have just explained, frequentist inference gives us a single point estimate 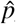 for the probability of connection, normally surrounded by a confidence interval which may be too large to be of any practical use and also, because it is flat, may give the illusion that the true value of *p* might be anywhere within this interval with equal probability. By contrast, Bayesian inference is more informative because it gives us a full probability density function *f_p_*(*p*), called the posterior, telling us exactly how likely every possible value of *p* actually is, given the collected data. In this way, even when confidence intervals overlap as is the case for practically all the SPN to SPN connections here (Figure 1A and B), which in a frequentist interpretation would lead us to dismiss the difference as non-significant without insight as to whether this is due to insufficient data or a true non-difference (Dienes, 2014; Makin and De Xivry, 2019), Bayesian inference gives us a much clearer picture of what the data can tell us. We introduce here a simple Bayesian approach to calculating the full posterior *f_p_*(*p*) for each type of connection from any pair-wise intracellular recording data.

As we show in the Methods, for these data, Bayesian inference turns out to be simple. Given the number of pairwise tests *n*, and the number of successful connections *k*, the posterior for *p* is a Beta distribution with updated shape parameters:

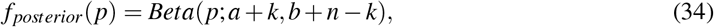

given initial values for its two parameters *a* and *b*. These initial values define the prior distribution for *p*, which reflects our initial beliefs about the possible values of *p*.

For instance, if we had initial values of *a* and *b* equal to 1, and we were looking at the data concerning D1 → D1 connections obtained by Taverna et al. (2008) (see Table 1) with *n* = 38 tests and *k* = 5 connections found, then we would obtain *a* = 6 and *b* = 34, and the resulting posterior would be the one depicted in Figure 1C (light blue curve). Depending on our assumptions, different values of *a* and *b* can be used to give the prior a desired shape. We begin with the common choice of the uniform distribution in which *p* could be anywhere between 0 and 1 with equal probability, achieved by setting *a* = *b* = 1 as in the example just given.

Using this prior in combination with the data of Taverna et al. (2008) gives us the posterior curves shown in Figure 1C. Once obtained, we can revert, if necessary, to a more frequentist standpoint by extracting from these density functions a single point estimate 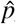, typically the Maximum A Posteriori value or MAP which is simply the value of *p* for which *f_p_*(*p*) is maximum, and a credibility interval around that MAP, which is the Bayesian equivalent of a confidence interval. The MAPs concentrate around relatively low values of *p* and their exact values, which are given in Table 2, lie between 0.06 for D1 → D2 pairs and 0.28 for D2 → D1 pairs. The uncertainty surrounding *p* is given by the width of the posteriors and of the 95% credibility intervals underneath the curves. For the data from Taverna et al. (2008), the connections with the smallest credibility interval of about 0.1 are the D1 → D2 pairs while the least well resolved connections are the D2 → D1 pairs for which the credibility interval spans roughly 0.2. By contrast, when we apply the uniform prior to the data of Planert et al. (2010), the D2 → D1 connections have the smallest uncertainty and the D2 → D2 connections the largest (Figure 1D).

**Table 2.** Maximum A Posteriori (MAP) estimates for the different probabilities of connection between SPNs, using different priors and different experimental studies. The “literature” prior is based on data on pairwise connections from intracellular recording studies that predated techniques for identifying types of SPN, as explained in the main text.

| Study | pair | uniform | Jeffreys | literature |
| --- | --- | --- | --- | --- |
| Taverna et al. (2008) | D1 → D1 | 0.132 | 0.122 | 0.116 |
| | D1 → D2 | 0.064 | 0.054 | 0.069 |
| | D2 → D1 | 0.277 | 0.272 | 0.222 |
| | D2 → D2 | 0.179 | 0.175 | 0.161 |
| Planert et al. (2010) | D1 → D1 | 0.070 | 0.060 | 0.074 |
| | D1 → D2 | 0.045 | 0.038 | 0.054 |
| | D2 → D1 | 0.125 | 0.120 | 0.117 |
| | D2 → D2 | 0.226 | 0.217 | 0.172 |

The fact that we used the same prior for all pairs of neuron types reflects our initial belief that there is no difference in the probability of connection between pairs. To overcome this belief requires a sufficient amount of evidence, and we can start comparing the different probabilities of connection visually by looking at how much the different posteriors overlap. Based on Figure 1C for example, it seems that in the data of Taverna et al. (2008) probabilities of connection segregate depending on the nature of the presynaptic neuron in the pair. The posteriors involving presynaptic D1 neurons overlap considerably with one another, and their region of highest density is lower than for presynaptic D2 neurons who also show great overlap, while there is much less overlap between pairs with different presynaptic neurons. This becomes even more obvious when looking at the 95% credibility intervals drawn underneath the curves which show more or less overlap depending on the nature of the presynaptic neuron: the credibility intervals for pairs with a presynaptic D1 neuron share an overlapping interval roughly covering probabilities of 0.05 to 0.15, while the overlapping interval for connections with a presynaptic D2 SPN ranges between probabilities of about 0.20 to 0.28. A similar pattern repeats itself in the data of Planert et al. (2010) (Figure 1D) although the exact values of the overlapping regions are shifted towards 0 compared to Taverna et al. (2008), an effect which is potentially due to maximum distance of sampling as explained later. This opens the possibility that there is indeed an asymmetry in terms of probability of connection that is dependent on the subtype of the presynaptic neuron, with no or little effect of the postsynaptic target subtype, something we will explore more thoroughly later.

One of the main advantages of Bayesian inference is that it forces researchers to be explicit about their priors and gives them the opportunity to choose appropriate ones. In order to illustrate this, we applied three further priors to the experimental data. Firstly, the so-called non-informative Jeffreys prior sets *a* = *b* = 1/2. An intuitive way of understanding this prior is to picture ourselves at the very beginning of the experiment, waiting for the result of the very first paired stimulation and recording. This test will either be successful or not, meaning that the shape of the prior should give most and equal weight to these two outcomes (inset of Figure 1E). Figures 1E and F show the posteriors that result from using this prior and we can see how they are practically identical to the posteriors obtained with a uniform prior. This was also the case when using the Haldane prior for which *a* and *b* equal 0 (not shown).

Our third prior is based on prior data, for Bayesian inference also provides us with a principled way of integrating previous knowledge into the prior. Earlier work (Taverna et al., 2004; Czubayko and Plenz, 2002; Koos et al., 2004) quantified the rate of lateral connections between SPNs without distinguishing SPN subtypes and concluded that lateral connections occurred at a rate of about 0.12. Using this information, we can design a beta distribution with a mean of 0.12 and an arbitrary variance of 0.005 (see Methods) which serves as our third and final prior shown in the inset of Figure 1G. Although the posteriors are more clearly different from the ones obtained with the uniform and Jeffreys prior they still look very similar. In fact, we find that the MAP values for a given type of connection (Table 2) are very close whatever the choice of prior, and the previous observation that the curves seem to segregate according to the subtype of the presynaptic neuron is valid in every case. On the other hand, because this prior is more informative than the two previous ones, uncertainty is reduced, as witnessed by the smaller credibility intervals. Given this robustness to the different priors, we shall henceforth exclusively use the prior based on previous literature when analysing connections between SPNs.

### D1 neurons make fewer connections than D2 neurons

We previously observed that D1 neurons seem to make fewer connections than D2 neurons without necessarily targeting one subtype over the other, based on how the posterior distributions appear to segregate by presynaptic subtype in Figure 1. We can go beyond this qualitative analysis by calculating a density function *f*_Δ_ for the difference between two probabilities of connection. For instance, if we’re interested in the difference in the probability of connection between D1 → D1 and D1 → D2 pairs (Figure 2A), using the posterior distributions *f*_*D*1→*D*1_(*p*) and *f*_*D*1→*D*2_(*p*), we can find the density function for Δ_(*D*1→*D*1)-(*D*1→*D*2)_ by:

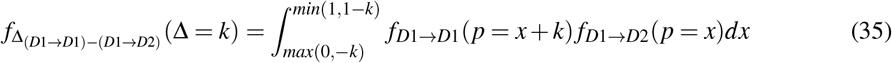

with the bounds of the integral such that *x* + *k* lies between 0 and 1. We can then calculate the probability that Δ_(*D*1→*D*1)–(*D*1→*D*2)_ is smaller than 0 by integrating this distribution between −1 and 0 (or calculate if it is greater than 0 by integrating the distribution between 0 and 1). By contrast, the frequentist strategy would be to compute a p-value giving the probability of getting an experimental result at least as extreme as the one observed assuming the null hypothesis of no difference in connection probabilities (i.e. Δ = 0), whereas the Bayesian approach allows us to calculate the probability that Δ is less than (or greater than) 0 given experimental results. Thus whereas the p-value tells us how surprising the actual data is if we accept the null hypothesis, the Bayesian approach can quantify precisely how unlikely the null hypothesis actually is.

**Figure 2.**
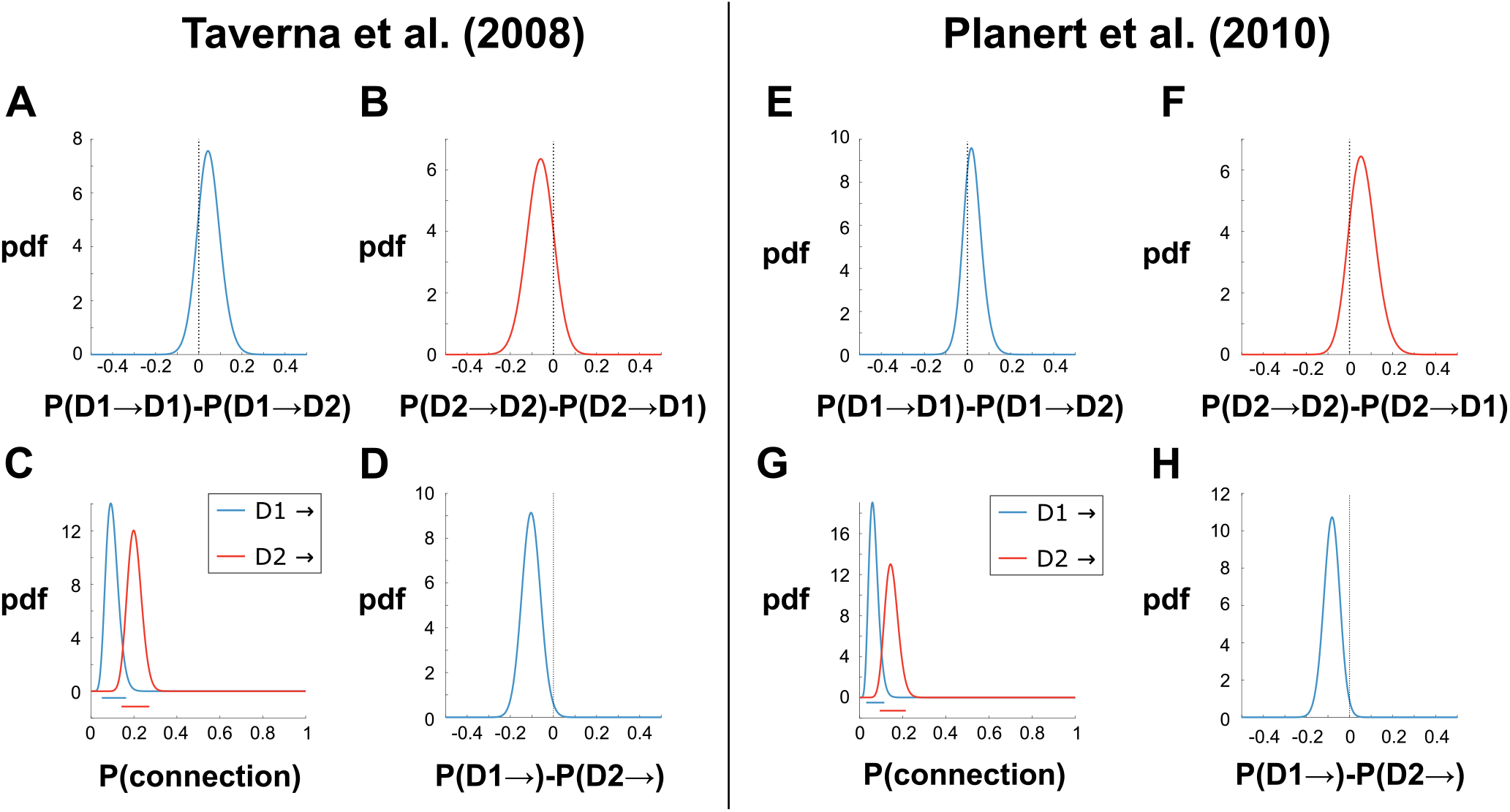
Comparison of the probabilities of connections between different SPN combinations. **A** Density function for the difference in the probabilities of connection in pairs with a presynaptic D1 neuron using data from Taverna et al. (2008). **B** Density function for the difference in the probabilities of connection in pairs with a presynaptic D2 neuron using data from Taverna et al. (2008). **C** Posterior density functions for the probabilities of connection collapsed according to the presynaptic neuron subtype. Bars underneath the curves correspond to the 95% credibility intervals. **D** Density function for the difference in connection probability between pairs with a presynaptic D1 neuron and pairs with a presynaptic D2 neuron. **E-H** Same as A-D, using data from Planert et al. (2010).

We applied this method to answer the question: do SPNs of either kind preferentially target SPNs of a certain subtype? In particular, based on their point estimates of connection probabilities, Taverna et al. (2008) have claimed that D1 neurons prefer to connect to other D1 neurons than to D2 neurons, a claim used in the computational study by Burke et al. (2017). As evidenced in Figures 2A and E which plot the density functions for Δ_(*D*1→*D*1)–(*D*1→*D*2)_ according to the data by Taverna et al. (2008) and Planert et al. (2010) respectively, this is not the case as both density functions include 0 among their most likely values. In fact, the probability that Δ_(*D*1→*D*1)–(*D*1→*D*2)_ is smaller than 0 is equal to 0.19 and 0.30 in Taverna et al. (2008) and Planert et al. (2010) respectively. Similarly, we do not find any difference when looking at whether D2 neurons have a preference for a particular postsynaptic neuron subtype (see Figures 2B and F for the density functions of Δ_(*D*2→*D*2)–(*D*2→*D*1)_): for Taverna et al. (2008), the probability that Δ_(*D*2→*D2*)-(*D*2→*D*1)_ is larger than 0 is 0.16, and the probability that it is smaller than 0 is 0.17 for Planert et al. (2010). We thus find no evidence that SPNs of one subtype (D1 or D2) preferentially target a certain subtype.

Having established that rates of connection are not different for a given presynaptic neuron subtype, we can collapse the data according to the subtype of the presynaptic neuron to answer another question: are D1 neurons more or less likely to make connections overall than D2 neurons? To do this, we simply add up the total number of tested pairs and connections found for the same presynaptic neuron type (e.g. for D1 SPNs: *n*_*D*1→*SPN*_ = *n*_*D*1→*D*1_ + *n*_*D*1→*D*2_ and similarly for *k*). In essence, this is equivalent to considering the posterior of one connection rate as the prior for connections with that same presynaptic neuron subtype, e.g. *f*_*D*1→*D*1_(*p*) is the prior for *f*_*D*1→*SPN*_(*p*). Figures 2C and G show the posterior distributions for the collapsed datasets, and both studies agree that D1 SPNs are less likely than D2 SPNs to make connections to other SPNs. The MAP values for connection rates from D1 neurons is 0.092 and 0.059 in Taverna et al. (2008) and Planert et al. (2010) respectively, versus 0.199 and 0.143 for connection rates from D2 neurons. If we look at the density function for the difference between the probability of connections for D1 and D2 neurons (Figure 2 D and H), the MAPs for *f*_Δ_(*D*1→*SPN*)–(*D*2→*SPN*)__ are −0.11 and −0.08 for Taverna et al. (2008) and Planert et al. (2010) respectively, while in both cases the probability that Δ_(*D*1→*SPN*)-(*D*2→*SPN*)_ is less than 0 is equal to 0.99. Thus, both studies contain very convincing evidence that D2 neurons are about twice as likely as D1 neurons to make connections to another SPN.

### Probability of connection as a function of distance

So far, when considering data on SPN connections from Taverna et al. (2008) and Planert et al. (2010) we have been careful to analyse each study separately, resisting the temptation of combining the two into a more powerful dataset. We were justified in being so careful since the two experiments used different maximum intersomatic distances between neurons, namely 50 *μm* in the study of Taverna et al. (2008) and 100 *μm* in that of Planert et al. (2010). Given that probability of connection between neurons typically decreases with distance (Hellwig, 2000; Humphries et al., 2010), sampling within a larger area around a neuron will probably cause a decrease in the ratio of connected pairs. Indeed, if we compare the probability of D1 or D2 neurons making lateral connections between the two experiments, we find that these probabilities tend to be larger in Taverna et al. (2008) which has the smaller sampling area, as illustrated by the corresponding density functions shown in Figure 3A. We thus introduce here a method for estimating the distance-dependent probability of connection between neurons types from data on neuron pairs recorded between some known maximum separation; our first use of this method is then to see whether it the distance-dependence is consistent between the two studies of SPN connectivity.

**Figure 3.**
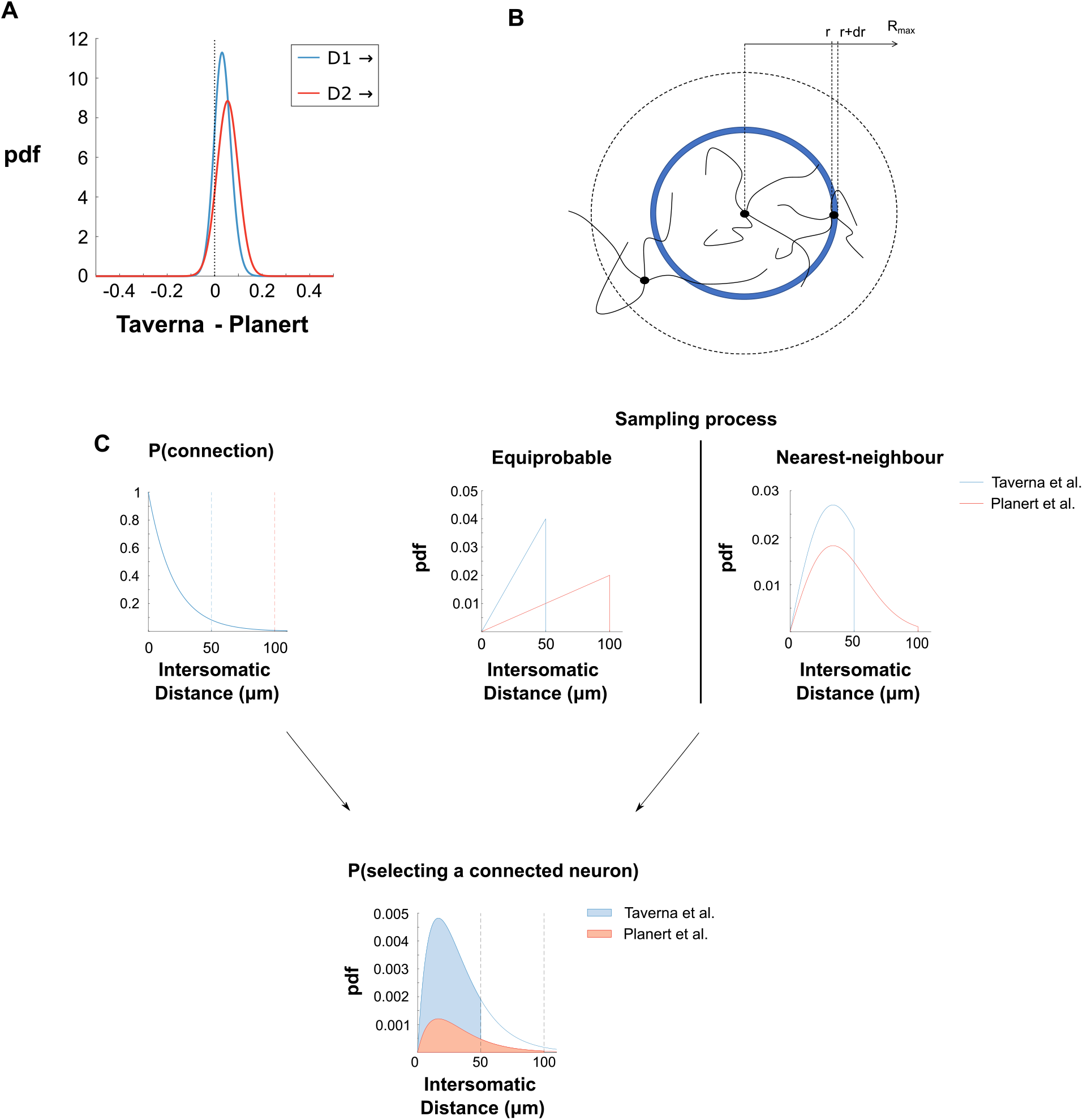
Estimating the probability of connection as a function of distance. **A** Density function for the difference in connection rates for a given presynaptic SPN type between the Taverna et al. and Planert et al. studies. **B** Experimenters chose their neurons within a certain maximum distance *R_max_* which defined a thin cylindrical volume of interest (here we draw the top of that cylinder). In the case of equiprobable sampling, the probability of choosing neurons further away increases as the infinitesimal volume corresponding to that distance increases as a linear function of *r*. **C** The probability of finding a connected pair of neurons depends on two different processes. Firstly, the process of connection, modelled by the probability of connection between two neurons given the distance between them, which we postulate decays exponentially; secondly, the process of sampling neurons in the experiment, modelled as the probability of selecting another neuron at a given distance from a starting neuron. We explore here two different scenarios for the sampling process: an equiprobable scenario in which neurons within a determined volume are selected randomly, and a nearest-neighbour scenario in which the selected neuron is whichever is the closest within the maximum distance set by the experimenters. The overall rate of connection reported by the experimenters then corresponds to the integral (shaded areas) of the product of these two probability models. Hence, differences in sampling processes can cause different rates of connection, even if the probability of connection given distance is the same.

To do this, we start by positing that this decrease obeys a simple exponential decay function:

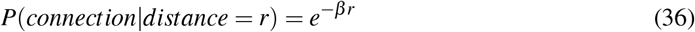

with *β* the decay parameter of unknown value, and *r* (for radius) the distance separating the two neurons. Ideally, to estimate this *β* parameter would require knowledge about the exact distance between every recorded pair of neurons, from which we could directly fit the model, but with simple assumptions on the sampling method used by experimenters, we can find an alternative way of converting values of *β* into *p*. Since the distance between each sampled pair of neurons in an experiment is in fact unknown to us, we shall consider it as a random variable. We can now express *p* as a function of *β* as

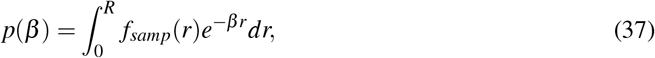

which is the product of the probability *f_samp_*(*r*) of experimenters selecting a neuron at distance *r* from another, and of the probability of these neurons being connected knowing *r* (equation 36) integrated over all possible values of *r* (see Figure 3 C for a visual depiction of what equation 37 means). *R* is the maximum distance at which the experimenters are sampling neurons, equal to 50 or 100 *μm* in Taverna et al. (2008) and Planert et al. (2010) respectively. We now need to find *f_samp_*(*r*).

A simple model for *f_samp_* would be that, given a certain volume surrounding a central neuron, the probability of sampling any given neuron in that volume is equiprobable for all neurons (Figure 3B) – we call this model *f_equi_*. With this assumption, we obtain the solution (see Methods):

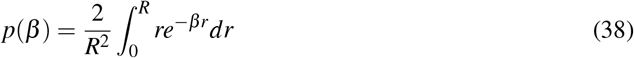

which gives us the corresponding value of *p* for any desired value of *β*. As we have posteriors *f_p_*(*p*) for the probability of connection between two types of neuron, we can now transform these into posteriors for *β*, *f_β_*(*β*) through parameter substitution using equation 38 (see Methods).

### Probability of connection decreases faster for D1 than for D2 neurons

We apply this method to the posteriors for the probabilities of connection collapsed according to the subtype of the presynaptic SPN and obtain the posteriors for the decay rate *β* shown in Figure 4A and B for D1 and D2 neurons respectively. Despite not being perfect, there is a good level of agreement between the two studies, which give estimates in the same ballpark. In both cases, although the posteriors do not overlap much, they do in fact lie quite close to each other providing us with a continuous, albeit broad, range of possible values. In the case of D1 neurons, the decay rate is expected to be in a region between 0.03 and 0.13 *μm*^-1^. The exponential decay curves representing the probability of connection as a function of distance for a decay rate equal to the MAP of each study (which are given in Table 3) are also shown in the inset of Figure 4A, and it is evident that they are extremely close to one another. As for D2 neurons (Figure 4B), the decay rate is smaller, as expected given that we have already shown that the overall probability of connection is higher for these neurons, ranging between 0.02 and 0.07 *μm*^-1^.

**Figure 4.**
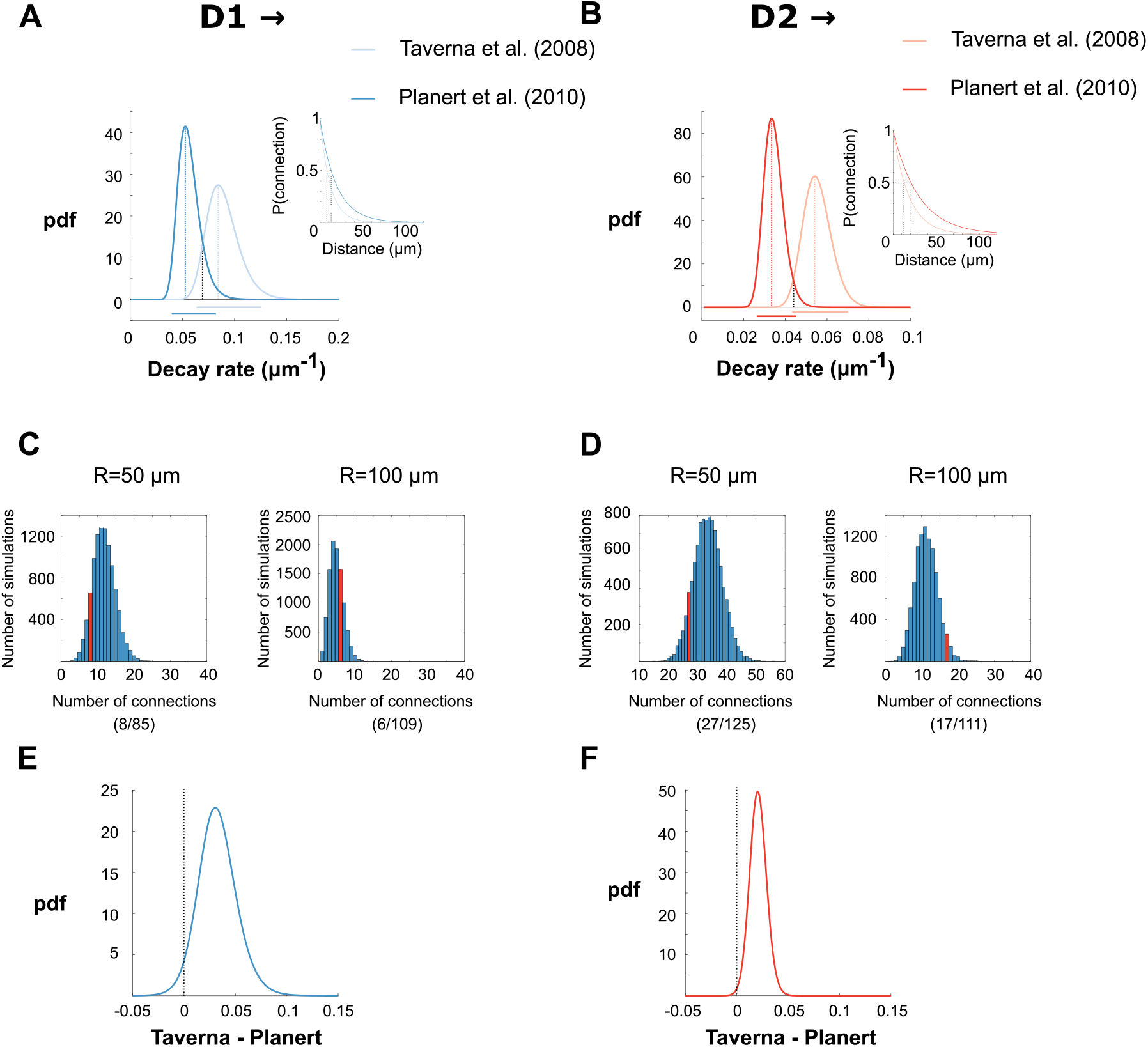
Estimates for the distance-dependence of connection probability between SPNs **A & B** Posterior density functions for the decay parameter of an exponential function representing the probability that a D1 or D2 neuron connects to a neighbouring neuron. Bars underneath represent 95% credibility intervals. Vertical black dashed line indicates the value of *β* at the maximum intersection of the two posteriors. Inset: Probabilities of connection given distance using the MAP values from the decay rate posteriors. **C & D** Monte Carlo simulations in which the best intersection estimate of *β* from **A & B** is used to try and replicate the exact experimental results of Taverna et al. (2008) (left) and Planert et al. (2010) (right) concerning pairs with a D1 presynaptic neuron (**C**) or a presynaptic D2 neuron (**D**). The exact results obtained by the experimenters correspond to the red bars and given between brackets underneath the bar graphs. **E & F** Density functions for the difference in decay rates between the two studies.

**Table 3.** MAPs and 95% credibility intervals (in *μm*^-1^) of the posterior curves for *β*.

| Study | Pair | $\hat{\beta}_{MAP} (\mu m^{-1})$ | 95% Credibility Interval |
| --- | --- | --- | --- |
| Taverna et al. (2008) | D1 SPN → SPN | 0.084 | [0.064, 0.125] |
| | D2 SPN → SPN | 0.054 | [0.043, 0.070] |
| Planert et al. (2010) | D1 SPN → SPN | 0.053 | [0.040, 0.082] |
| | D2 SPN → SPN | 0.033 | [0.026, 0.045] |
| | FSI → D1 SPN | 0.002 | [0.0004, 0.009] |
| | FSI → D2 SPN | 0.006 | [0.002, 0.017] |
| Gittis et al. (2010) | FSI → D1 SPN | 0.004 | [0.003, 0.005] |
| | FSI → D2 SPN | 0.007 | [0.005, 0.009] |
| | FSI → FSI | 0.003 | [0.001, 0.008] |
| Ibáñez-Sandoval et al. (2011) | NGF → SPN | 0.002 | [0.001, 0.006] |

To get a better idea of how consistent the results are between the Taverna et al. (2008) and Planert et al. (2010) datasets, we used Monte Carlo simulations to try and recover the number of observations made by each experiment with its own value of *R* using the *β* value that seems most likely given the two posterior curves, i.e. the intersection of the two posterior curves (black dotted lines in Figure 4A,B). We ran 10000 virtual experiments by generating random distances between pairs of neurons (according to equation 14 in the Methods) and the maximum distance *R* used by that study, and then deciding whether they were in fact connected according to the probability of connection of equation 36. We generated the same number of pairs as tested in each study and then reported the number of times we obtained the exact same number of positive results (red bars in the histograms of Figure 4C and D). For instance, setting an intermediate decay rate of 0.075 *μm*^-1^ for D1 neurons, and generating 10000 replications of the experiment of Taverna et al. (2008) which recorded 85 pairs of SPNs with a D1 presynaptic neuron, we obtained more than 500 simulations where exactly 8 pairs were connected, which is the number originally reported (Figure 4C left figure). In all four cases, we managed to replicate the original results relatively often, proving that the best estimates for *β* are reasonable. As a sanity check, we also used the decay rate MAPs for one subtype to try and replicate the results of the other subtype and found it much harder if not impossible to replicate the results, verifying that the decay rates have to be different for the two types of SPNs (results not shown).

If we compare the estimates between the two datasets more critically, there is a clear bias for the posterior curves extracted from Taverna et al. (2008) to be shifted to the right compared to the posterior curves from Planert et al. (2010) (Figure 4A and B). Indeed, if we refer to the insets in Figure 4A and B, the best estimate of *β* according to the study by Taverna et al. (2008) would predict a 50% drop in probability of connection every 8 *μm* for D1 neurons versus every 13 *μm* according to the study by Planert et al. (2010). In the case of D2 neurons, the difference between the two exponential curves is even greater, with a half distance of 13*μm* versus 21 *μm* according to Taverna et al. (2008) and Planert et al. (2010) respectively. Furthermore, the density functions for the difference in posteriors between the two studies both lie predominantly in the positive domain (Figure 4E and F) and the probability that the decay rate is larger in the Taverna than Planert data is 0.967 and 0.996 for *β*_*D*1→*SPN*_ and *β*_*D*2→*SPN*_ respectively. Though the disagreement between the data-sets is small, it is nonetheless consistent.

### Biased neuron sampling could explain differences between datasets

One potential explanation is that the sampling of neuron pairs was more complex than the equiprobable sampling model we first assumed. In this section, we explore this possibility by considering a model for *f_samp_* where, rather than choose neurons equiprobably within a visible area surrounding a neuron, experimenters preferentially tested neurons which were closer to one another to maximise the probability of detecting connections.

To explore this model, we used the same probability of connection given distance (Equation 36) in combination with a new density function *f_NN_* for the probability of the distance to the nearest neighbour, to derive a new mapping from *p* to *β*. We found the resulting mapping to be (Methods):

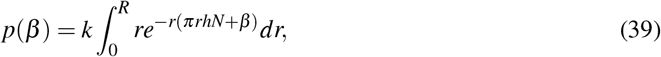

where *k* is a normalising constant. Contrary to the previous equiprobable sampling model (Equation 38), where these parameters cancelled out, this mapping depends on *N*, the density of SPNs in the striatum, and *h*, the height of the cylinder in which sampling takes place. We used here an estimate of the SPN density in mice of 80500 per mm^3^, and tested three different values of *h* to get three different nearest-neighbour distributions: 0.1 *μm*, 1 *μm* and 10 *μm*. We then use this mapping to transform the posteriors for *p* into posteriors for *β* as before (Methods).

The first column of Figure 5 shows the resulting posteriors for D1 neurons. The picture for *h* = 0.1*μm* is not so different from that obtained under the equiprobable sampling hypothesis, but for greater values of *h*, the posteriors overlap far more. In particular, for *h* equal to 1 *μm* the posterior curves practically coincide. Similarly for D2 neurons, *h* = 0.1*μm* does not much improve the agreement between the two studies, but greater values of *h* do (Figure 5, second column). This approach successfully illustrates how a tendency to select neurons closer together might account for the discrepancy observed in estimates of the decay parameter using the simpler equiprobable sampling model. Moreover, this analysis shows how the details of data sampling matter when estimating connectivity statistics from intracellular recording data.

**Figure 5.**
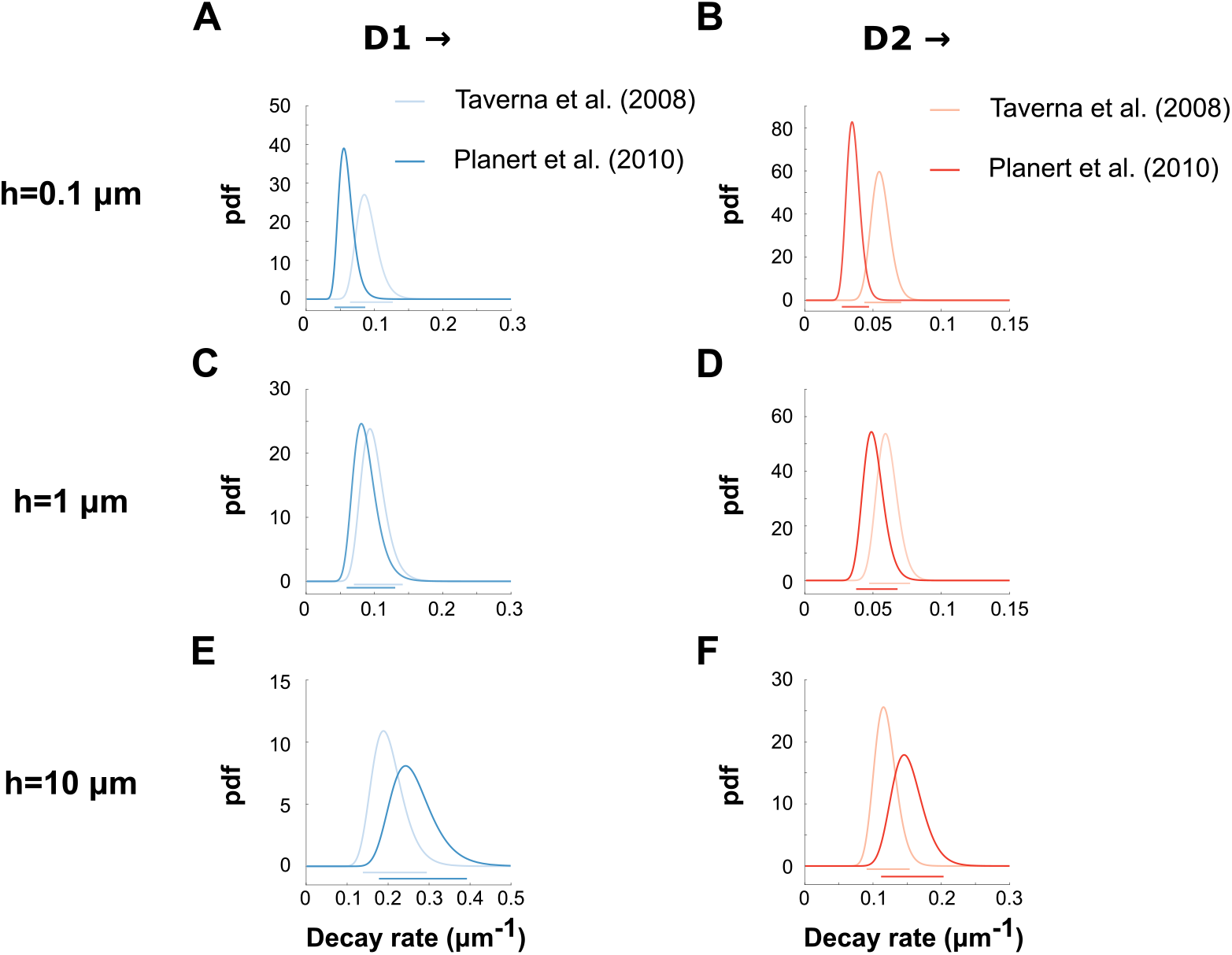
Density functions for the decay parameter assuming a nearest-neighbour model of neuron selection for different values of the depth *h* of the sampling region. **A** Density function for *β* for D1 neurons when *h* = 0.1*μm*. **B** Density function for *β* for D2 neurons when *h* = 0.1*μm*. **C-D** Same as **A-B** for *h* = 1*μm*. **E-F** Same as **A-B** for *h* = 10*μm*.

### Fast Spiking interneurons preferentially connect to D1 SPNs

We now turn to completing our Bayesian map of the striatal microcircuit by evaluating the connections of different species of interneurons to the SPNs and to each other - we present the full map in the Discussion (Figure 10). Three main types of interneurons are commonly documented (Kreitzer, 2009): Fast Spiking (FS), Persistent Low Threshold Spiking (PLTS), and cholinergic (Ach) interneurons. We will also include in this list tyrosine-hydroxylase (TH) and NPY-NGF interneurons because of their relationship to Ach interneurons, which is crucial to the function of the cholinergic component of the circuitry (Ibáñez-Sandoval et al., 2011; English et al., 2012; Dorst et al., 2020). We took data on pairwise intracellular recordings of these interneuron types from the range of studies listed in Table 1, and determined Bayesian posteriors for *p* as previously. Unlike for the SPNs we did not have prior studies of the interneuron connections to help us design a prior, so we relied on a uniform prior instead.

We focus first on FS interneurons that project to the SPNs, using intracellular recording data from Planert et al. (2010) and Gittis et al. (2010). The posteriors we obtain for Planert et al. (2010) (Figure 6B) are consistent with quite high connection probabilities: the MAPs are 0.67 and 0.89 for connections to D2 and D1 neurons respectively. However, the small size of the samples means that the range of possible values is also broad (95% credibility intervals: D2, [0.35, 0.88]; D1, [0.56, 0.98]). Fortunately, the study of Gittis et al. (2010) is based on a much larger sample resulting in narrower posteriors shown in Figure 6A. Thanks to these narrower posterior curves, it is possible to conclude that FS interneurons preferentially target D1 neurons. In fact, when we integrate Δ_*FS*→*D*1-*FS*→*D*2_ (Figure 6A inset), we find the probability that FS interneurons prefer to connect to D1 neurons is greater than 0.99.

**Figure 6.**
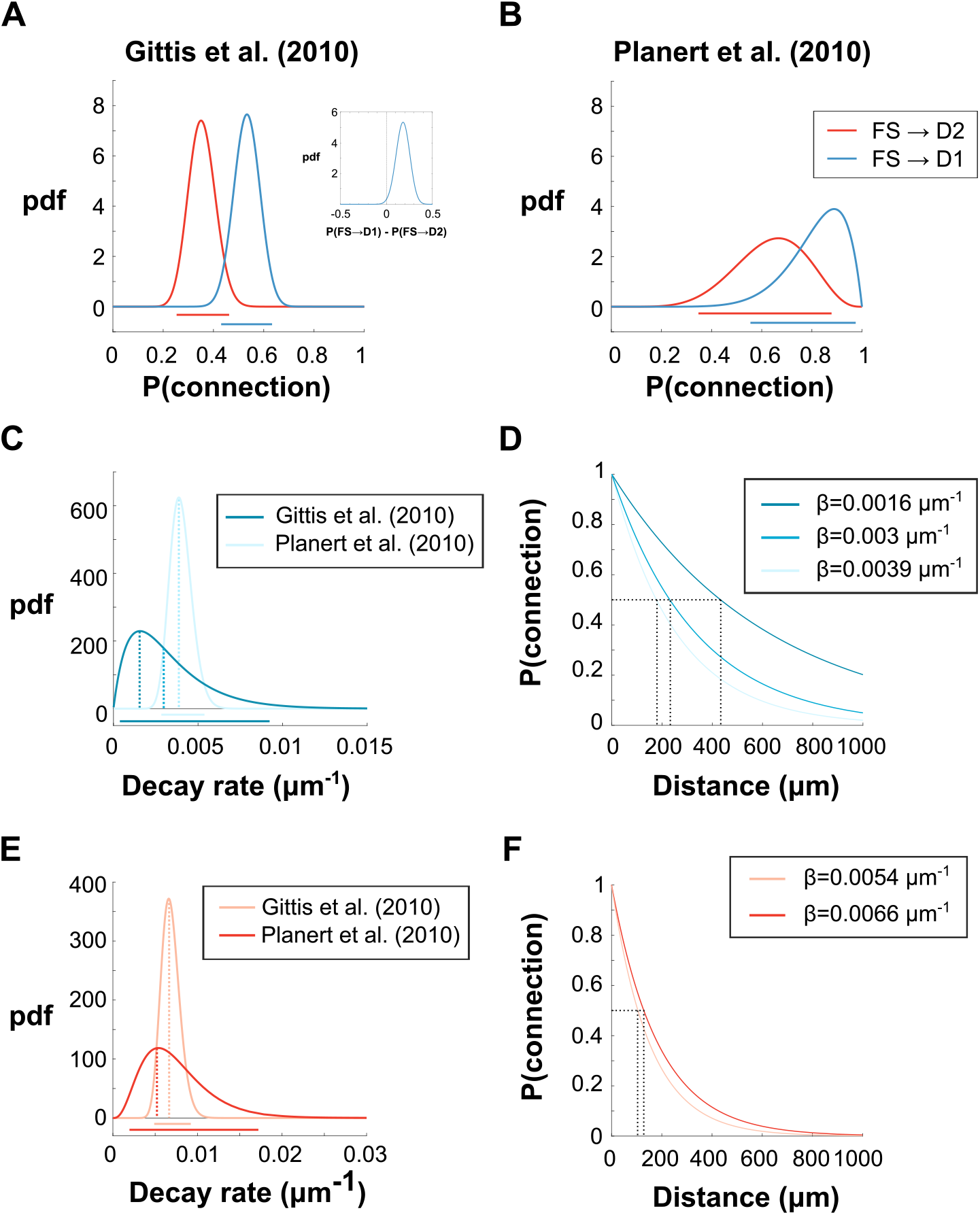
Bayesian analysis of connection probabilities of FS interneurons onto D1 and D2 SPNs. **A** Connection probabilities of fast spiking (FS) interneurons connecting to D1 and D2 SPNs according to the data of Gittis et al. (2010) who set a maximum distance of 250 *μm* between neurons. Inset: density function for the difference in probability of connection. **B** As for panel A, according to the data of Planert et al. (2010) who used a maximum distance of 100 *μm* instead. **C** Posterior density functions for the decay rate of probability of connection for FS→D1 pairs assuming equiprobable sampling of neurons. **D** Probabilities of connection given distance for three different values of *β* corresponding to the MAP estimates of each study and the intersection of the two posterior curves. **E-F** Same for FS→D2 pairs as C and D respectively. Because the MAP estimate for Planert et al. (2010) coincides with the intersection of the two posteriors, only two exponential decays are tested in F.

The two studies seem to disagree as Planert et al. (2010) gives much higher estimates of *p*, but this is resolved by taking into account the maximum distance used by the two studies, 100 *μm* for Planert et al. (2010) and 250 *μm* for Gittis et al. (2010). Indeed, if we convert the posteriors for *p* into posteriors for the exponential decay rate *β* of the probability of connection given distance, with the assumption of equiprobable sampling as previously explained, we obtain posteriors which very largely overlap (Figure 6C and E; Table 3) thus reconciling the two studies. In line with the already discussed overall smaller rate of connection to D2 neurons, the probability of connection drops much faster as distance increases for connections to D2 neurons (dropping to 50% after about 100 *μm*, see Figure 6F) than for connections to D1 neurons (50% connection rates occurring at a distance of at least 200 *μm*, see Figure 6D). Consequently, there are at least two length-scales in the striatal microcircuit, with connections between SPNs falling to 50% probability within a few tens of micrometers (Figure 4A,B), but connections to SPNs from FS interneurons falling to 50% probability at a hundred micrometers or more (Figure 6D,F).

### PLTS interneurons make few local connections in striatum

We turn now to the connections that FS interneurons make on other interneurons of the striatum. To assess these, we analysed data from Gittis et al. (2010) on connections FS interneurons make to PLTS, cholinergic, and other FS interneurons (Figure 7A). Their data on connections between FS and PLTS interneurons were pooled with the data on the same connections from Szydlowski et al. (2013): We checked that the data from the two studies were in agreement by calculating posteriors separately for each study and found the density functions for the difference between the posteriors (Δ) for both directions (FS→PLTS and PLTS→FS) included 0 among their most likely values (results not shown).

**Figure 7.**
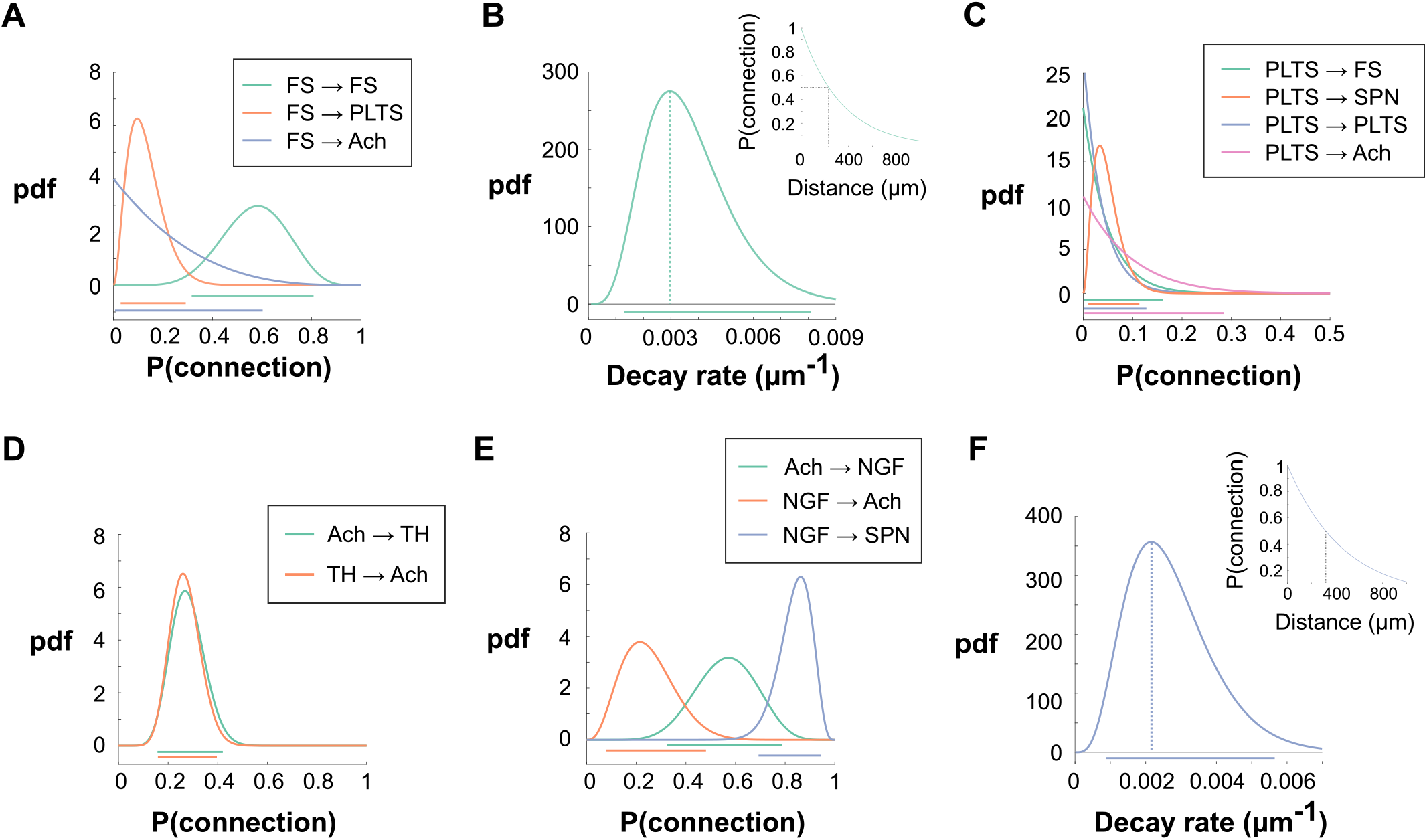
Bayesian analysis of connection probabilities between striatal interneurons using a uniform prior. **A** Posterior density functions for FS interneurons connections onto other interneurons according to the data of Gittis et al. (2010). **B** Posterior density functions for the decay rate of probability of connection for FS → FS pairs assuming equiprobable sampling of neurons. Inset: Exponential decay function for the probability of connection between pairs of FS interneurons corresponding to the MAP estimate of the decay rate. **C** Posterior density functions for PLTS interneuron connections onto other interneurons according to the data of Gittis et al. (2010). **D** Posterior density functions for connections between cholinergic and TH interneurons according to data from Dorst et al. (2020). **E** Posterior density functions for connections between cholinergic interneurons, NGF interneurons and SPNs according to the data of English et al. (2012) and Ibáñez-Sandoval et al. (2011). **F** Posterior density functions for the decay rate of the probability of connection for NGF → SPN pairs assuming equiprobable sampling of neurons. Inset: Exponential decay function for the probability of connection between NGF → SPN pairs corresponding to the MAP estimate of the decay rate in E.

We see from these data that the probability of connection from FS interneurons to PLTS interneurons is low 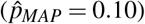 but uncertainty regarding these connections is quite large (95% credibility interval = [0.03, 0.29]), while the probability of connection to cholinergic interneurons is even more uncertain with a credibility interval ranging from 0 to 0.60 and therefore requires more investigation. Connections between FS interneurons are relatively common 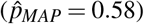 but with broad uncertainty (95% credibility interval = [0.32, 0.81]). While this broad uncertainty in *p* translates into a broad uncertainty for the decay rate *β* of the probability of connection given the distance between a pair of FS interneurons (Figure 7B), we see that the distance-dependence for pairs of FS interneurons is similar to that for connections of FS interneurons to SPNs with a half-distance for connection probability as a function of distance of about 200 *μm* for 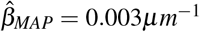. Unfortunately, Szydlowski et al. (2013) do not provide a maximum distance which prevents us from transforming *f_p_*(*p*) for FS → PLTS pairs into the corresponding *f_β_*(*β*).

The combined data of Gittis et al. (2010) and Szydlowski et al. (2013) (Figure 7C) shows no evidence that PLTS interneurons connect locally to other interneurons (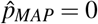 for all pairs), but connections to SPNs, although sparse, are clearly established within a maximum distance of 250 *μm* (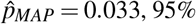 credibility interval = [0.010, 0.114]). Compared to FS interneurons, there is less uncertainty concerning the rates of connection for these PLTS interneurons, as evidenced by the smaller credibility intervals shown in Figure 7C. It remains to be seen whether there is an asymmetry in connection probability to D1 and D2 SPNs as the experimenters did not make this distinction when testing PLTS → SPN connections.

### Further evidence that the effect of cholinergic interneurons onto SPNs is mediated by GABA interneurons

A recent study (Dorst et al., 2020) reported intracellular recording data on connections between cholinergic interneurons and the subtype of GABAergic interneurons that express tyrosyne-hydroxylase (TH interneurons) (see Table 1). When we apply our Bayesian method to this dataset (Figure 7D), we find that TH and cholinergic interneurons connect reciprocally to one another quite frequently and with practically equal probabilities 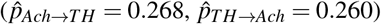, with uncertainty estimates which are relatively good compared to other interneuron connections (95% credibility interval = [0.159, 0.396] for TH → Ach connections, and [0.157, 0.420] for Ach → TH connections).

The activity of cholinergic interneurons indirectly affects SPNs via at least one type of GABAergic interneuron (English et al., 2012). To examine this route, we combine pairwise intracellular recording data from English et al. (2012) on connections between cholinergic interneurons and NPY-NGF interneurons, with data from Ibáñez-Sandoval et al. (2011) on connections from NPY-NGF interneurons to SPNs. Our analysis (Figure 7E), reveals that cholinergic neurons connect frequently to NPY-NGF interneurons, which in turn connect very frequently to SPNs, making them an effective relay of cholinergic signals; this relay may also be regulated by the NPY-NGF interneurons frequent feedback connections on cholinergic interneurons. Furthermore, given that Ibáñez-Sandoval et al. (2011) used a maximum distance between neurons of 100 *μm*, it is possible to transform the posteriors for the probability of NGF → SPN into posteriors for the exponential decay rate *β* of probability of connection given distance. The MAP of *β* is the lowest we have found at about 0.002 *μm*^-1^ (Figure 7F; Table 3). This means that even at a distance of 100 *μm* from an NGF interneuron, an SPN still has a 0.8 probability of receiving a connection from this interneuron, which partly explains the effectiveness of the cholinergic system in regulating SPN activity.

### Evidence used to compare SPN sub-type connection rates in wild type and Huntington’s disease mice is insufficient

To this point we have used our Bayesian approach to evaluate the probability of connection, the evidence for it, and (where possible) its dependence on the distance between neurons for every unique connection within the striatal microcircuit (for which there is extant data). We turn now to showing how our Bayesian approach lets us not just construct a map of the microcircuit, but also quantitatively test evidence for changes in the microcircuit. To do so, in this section we evaluate evidence that connections between SPNs change in a mouse model of Huntington’s disease (Cepeda et al., 2013); in the following section we evaluate evidence for how connections between SPNs change over development.

The study of Cepeda et al. (2013) used smaller samples of identified SPN pairs than the (Taverna et al., 2008) and (Planert et al., 2010) studies of SPN connectivity (see Table 4) and consequently the resulting posteriors are notably impacted by the choice of the prior (Figure 8A and D). Indeed, the posterior curves obtained from the data of Cepeda et al. (2013) with a uniform prior or the prior based on previous literature look very different, contrary to those of Taverna et al. (2008) (compare Figures 1C and G) and Planert et al. (2010) (compare Figures 1G and H). In particular, the posteriors for wild-type mice obtained from the prior based on the previous literature look very similar to the initial prior (Figure 8D) simply because of the small number of samples. Independently of the choice of prior, the posteriors overlap so much that it is not possible to confirm the rates of connection differ between any pairs of SPNs in the wild type mice (Figures 8A and D). Given how broad the posteriors are, we infer that this lack of difference is due to insufficient data rather than a true absence of difference.

**Table 4.** Experimental data from wild type and HD model mice from Cepeda et al. (2013), alongside results of the Bayesian analysis using either a uniform or literature prior. For each type of connection, *k* is the number of connections that were found and *n* the total number of tested connections. HD: Huntington’s disease.

| Data | Pair | $k$ | $n$ | Prior | $\hat{p}_{MAP}$ | 95% credibility interval |
| --- | --- | --- | --- | --- | --- | --- |
| Wild type | D1 SPN → D1 SPN | 4 | 14 | Uniform | 0.286 | [0.118, 0.551] |
|  |  |  |  | Literature | 0.170 | [0.079, 0.333] |
| | D1 SPN → D2 SPN | 2 | 10 | Uniform | 0.200 | [0.060, 0.518] |
|  |  |  |  | Literature | 0.124 | [0.048, 0.292] |
| | D2 SPN → D1 SPN | 2 | 10 | Uniform | 0.200 | [0.060, 0.518] |
|  |  |  |  | Literature | 0.124 | [0.048, 0.292] |
| | D2 SPN → D2 SPN | 4 | 14 | Uniform | 0.143 | [0.043, 0.405] |
|  |  |  |  | Literature | 0.109 | [0.042, 0.260] |
| HD model | D1 SPN → D1 SPN | 7 | 22 | Uniform | 0.318 | [0.164, 0.529] |
|  |  |  |  | Literature | 0.210 | [0.14, 0.359] |
| | D1 SPN → D2 SPN | 1 | 6 | Uniform | 0.167 | [0.037, 0.579] |
|  |  |  |  | Literature | 0.104 | [0.035, 0.283] |
| | D2 SPN → D1 SPN | 1 | 6 | Uniform | 0.167 | [0.037, 0.579] |
|  |  |  |  | Literature | 0.104 | [0.035, 0.283] |
| | D2 SPN → D2 SPN | 0 | 14 | Uniform | 0 | [0, 0.218] |
|  |  |  |  | Literature | 0.048 | [0.013, 0.180] |

**Figure 8.**
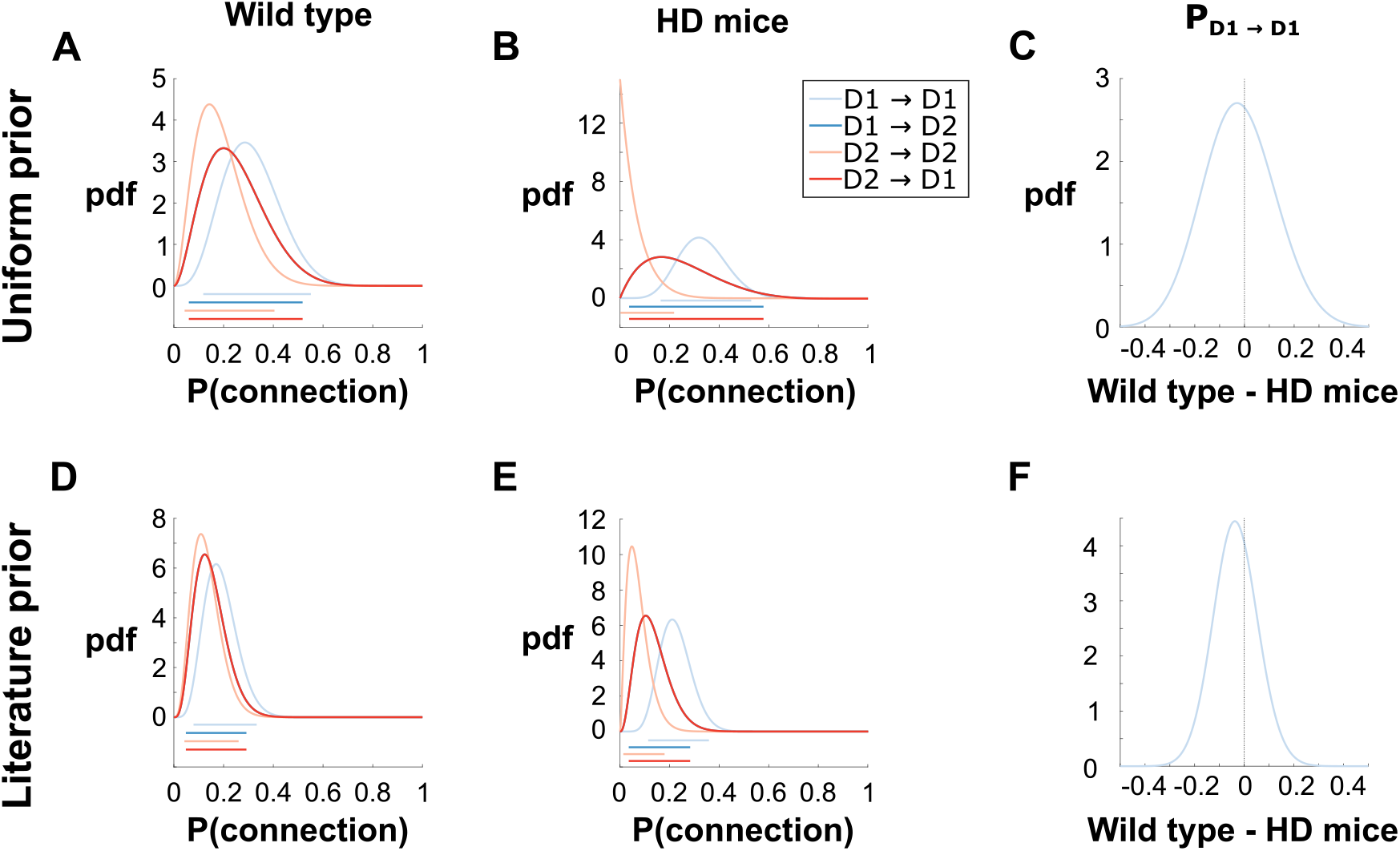
Bayesian analysis of the study by Cepeda et al. (2013), comparing the probability of lateral SPN connections in wild type mice and a model of Huntington’s disease. **A** Posterior density functions for the probabilities of connection in the wild type mice using a uniform prior. Bars underneath represent the 95% credibility intervals. The curves for D2 → D1 and D1 → D2 coincide exactly. **B** Posterior density functions for Huntington’s disease animals using a uniform prior. The curves for D2 → D1 and D1 → D2 also coincide exactly. **C** Probability density function for the difference in probabilities of connection for D1 → D1 pairs between the two animal groups. **D-F** Same as **A-C** using the prior based on the past literature.

Crucially, this lack of data is also true when comparing connection rates between the wild type and Huntington’s model mice (Figure 8A-B or D-E). We find no evidence to support one of the conclusions reached by the authors that D1 → D1 connections are more likely in the Huntington’s model than wild-type mice. Indeed, if we plot the density functions for the difference in probabilities of connection for D1 → D1 pairs between the two animal groups, we can see that it is very probable for this difference to be 0, but also any other value ranging from roughly −0.3 to 0.3 if we use a uniform prior (Figure 8C) or a slightly more conservative −0.2 to 0.15 using the prior based on previous literature (Figure 8F).

### D2 SPN connection asymmetries appear during development

The development of connections between SPNs and their asymmetry can be tracked through postnatal development thanks to a recent study by Krajeski et al. (2019) who measured the probability of connection for different SPN pairs at three different stages of post-natal mouse development. The researchers reported that D1 neurons established lateral connections earlier than D2 neurons (a reproduction of these results with our added Wilson confidence intervals is shown in Figure 9A-C). We used our Bayesian approach to check this conclusion against the uncertainty in the experimental data, and provide further details of the development of the striatal microcircuit.

**Figure 9.**
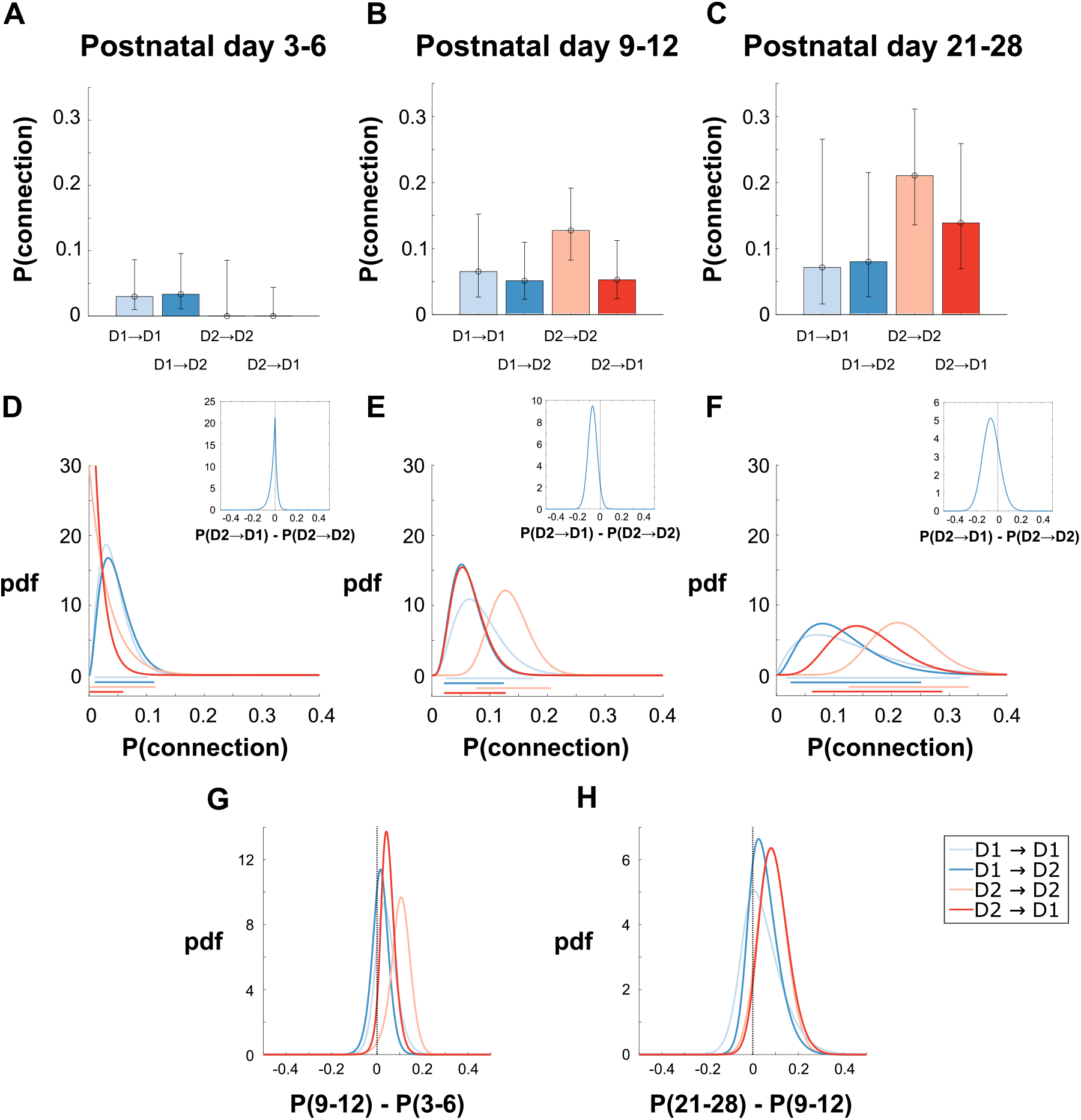
Post-natal development of the lateral connections of SPNs using data from Krajeski et al. (2019). **A-C** Point estimates of the probabilities of connection at different developmental stages from Krajeski et al. (2019). We add here the 95% Wilson confidence intervals. **D-F** Posterior probability density functions for the probability of connections between SPNs at each developmental stage. Coloured bars underneath the plot represent the 95% credibility intervals. A uniform prior as in Figure 1C is used. Inset: Density function for the difference in probability of connection for pairs with a D2 presynaptic neuron. **G-H** Density functions for the difference in connection probabilities for each pair of neuron types between consecutive stages of postnatal development.

In order to apply our Bayesian method to this data we resort to the uniform prior, since we have no particular expectation about these connection rates at these stages of development, and obtain the posteriors for each combination of neurons shown in Figure 9D-F. We have also tested the Jeffreys prior and obtained practically identical results (not shown). We can see that D1 neurons have already made some connections in the first 3-6 days of postnatal development (P3-6; Figure 9D), but it is hard to say from inspecting the posteriors at each subsequent developmental stage (P9-12 and P21-28) whether the connections made by D1 SPNs continue to develop or have already finished by P3-6 (Figure 9E and F). If we instead look at the difference in posteriors (*f*_Δ_) between consecutive developmental stages (Figure 9 G and H), we see some evidence that connections made by D1 neurons continue to develop, with respective probabilities of 0.81 (P3-6 to P9-12) and 0.77 (P9-12 to P21-28) that the connection density of D1 neurons increases (probabilities again found by integrating *f*_Δ_ between 0 and 1). Comparing the earliest (P3-6) and latest (P21-28) stages gave a similar probability of 0.85 that D1 connections increased (not shown).

For presynaptic D2 neurons on the other hand, it is clear that no or very few connections are present at P3-6, and that they gradually appear later (Figure 9D-F). Computing the difference in posteriors (*f*_Δ_) for *P*(*D*2 → *D*2) and *P*(*D*2 → *D*1) between consecutive stages (Figure 9 G and H), we find the probability that D2 SPNs increase their connection density is 0.95 between P3-6 and P9-12 and 0.92 between P9-12 and P21-28, indicating a gradual development of these connections up to postnatal days 21-28. By also calculating *f*_Δ((*D*2→*D*1)-(*D*2→*D*2))_ at each of these three stages (plotted as insets in Figure 9 D-F), we find good evidence that D2 neurons first connect to other D2 neurons before connecting to D1 neurons, supporting the claim by the authors of the original article (Krajeski et al., 2019). Notably, our analyses in this paper have thus shown that while there is no evidence for a difference in the probability of pre-synaptic D2 SPNs connecting to D1 or D2 SPNs in the adult striatum (Figure 2B,F), there is evidence that the D2 → D1 and D2 → D2 connections develop at different rates.

Our finding of strong evidence that D1 neurons are less likely to receive connections from SPNs than D2 neurons, both in adults (Figure 2) and at later stages of development (Figure 9), implies a role for active wiring processes in the developing striatum. We considered a simple model of a passive wiring process in which the contact of an axon from a first neuron onto the dendrite of a second neuron is determined only by the probability that an axon segment and a dendritic segment simultaneously occupy the same location (Liley and Wright, 1994; Kalisman et al., 2003; Humphries et al., 2010). For SPNs we have the repeated observation that D1 SPNs have denser dendritic trees for the same volume as D2 SPNs (Gertler et al., 2008; Fujiyama et al., 2011; Gagnon et al., 2017). A passive wiring model would thus predict that D1 SPN dendrites receive more axonal contacts than D2 SPN dendrites from the same pre-synaptic type of SPN.

If this were true, we would expect to find in our analyses here that the asymmetry of connection rates would depend on the type of the postsynaptic neuron, when we instead find it depends on the type of presynaptic neuron; and we would expect D1 → D1 connections to be quite numerous when in fact these are quite rare. Together with our confirmation that the data of Krajeski et al. (2019) show D1 and D2 neurons develop their connections at a different rate, our analyses thus suggest that there is an active wiring process in striatal development that causes either an underexpression of connections to D1 SPNs, or overexpression of connections to D2 SPNs.

## DISCUSSION

We presented a Bayesian inference approach to analysing connectivity using intracellular recording data, and applied it to reconstruct the microcircuit of the striatum from an exhaustive survey of data from pairwise intracellular recordings. None of these data have had any assessment of the uncertainty in their connection estimates or of the strength of evidence they provide. Our new approach allows us to now draw rigorous conclusions about the strength of evidence for claims about the microcircuit, and in turn synthesise these data into as complete a map as the data allow.

### A Bayesian map of the striatal microcircuit in mice

Figure 10 synthesises the complete map we obtained of the striatal microcircuit in mice. It emphasises our key results: first, there is strong evidence of a connection asymmetry that depends on the type of presynaptic SPN – namely that D2 SPNs are roughly twice as likely to contact another SPN as D1 SPNs – but no evidence of an asymmetry that depends on the type of postsynaptic SPNs; second, that there is strong evidence for FS interneurons preferentially connecting to D1 SPNs; third that there is strong evidence of dense projections from NPY-NGF interneurons to SPNs, likely as dense or denser than those from FS interneurons; and, finally, that connections between SPNs occur on much shorter length-scales than the connections made by interneurons.

**Figure 10.**
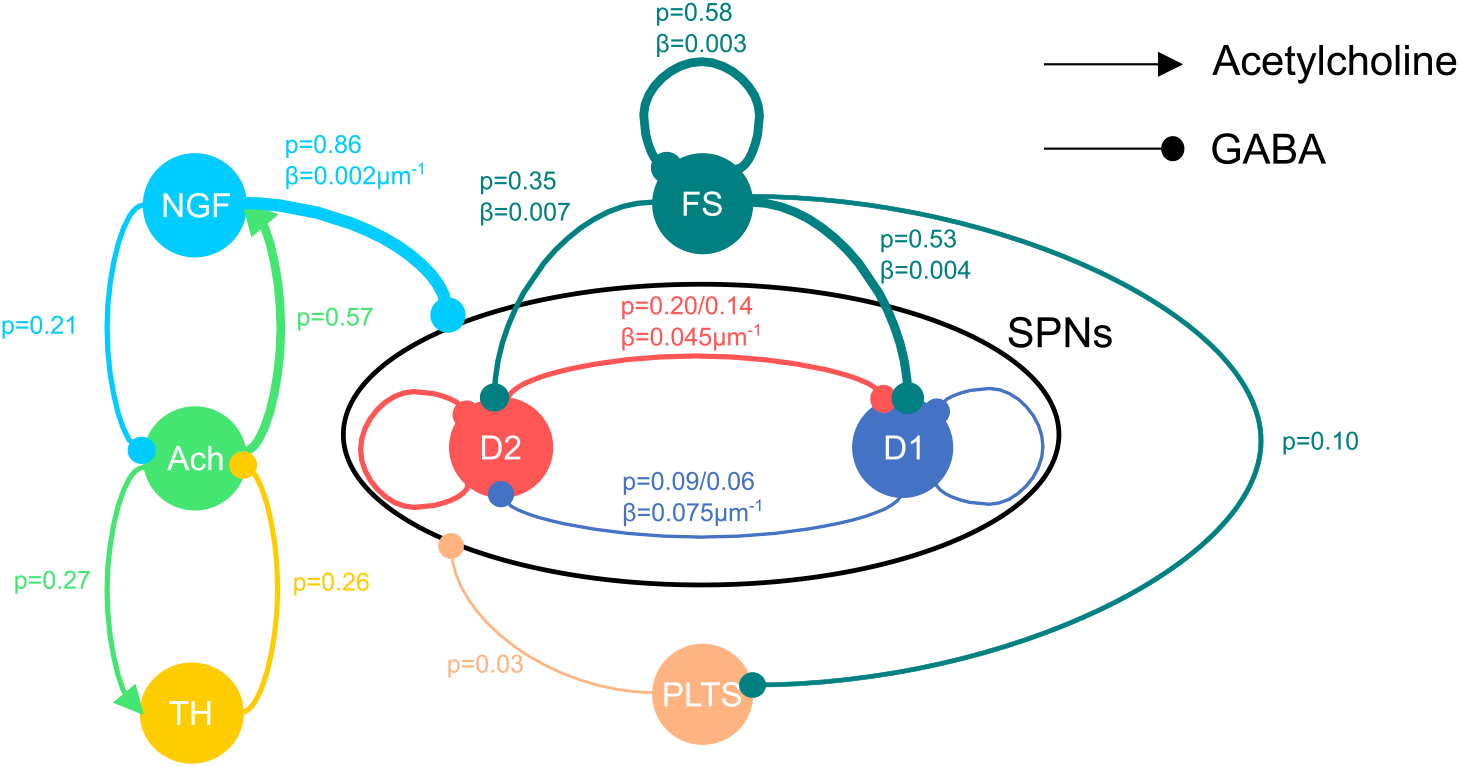
Map of the striatum microcircuitry based on the MAP estimates for *p* and, when a maximum intersomatic distance was available, the decay rate *β* assuming equiprobable sampling. Line thickness is indicative of the relative probability of these connections. Connections between and within SPN subtypes are assumed to be the same for a given presynaptic subtype, as established in the main text, and the two different estimates for *p* correspond to the two different maximum distances used in Taverna et al. (2008) and Planert et al. (2010). Modellers wishing to use this map should beware of the relative population size of these different neurons. For instance, although the probability of connection between SPNs is relatively small compared to connections from FS interneurons, this is potentially counterbalanced by the much greater number of SPNs within a given volume (Humphries et al., 2010). The map also necessarily omits known connections for which there are no appropriate intracellular recording data.

For ease of interpretation, Figure 10 summarises each connection probability as the best point estimate we can obtain (namely, the MAP of the corresponding posterior distribution). But of course we now have the full posterior distributions underlying these estimates, and some of these are broad – for example, the connection probability from NGF to Ach interneurons has a 95% credible interval twice as wide as its best (MAP) estimate (Table 1). Our posterior distributions thus reveal that a wide range of potential striatal wirings are consistent with current data.

From this it follows that any model of the striatum should sample its connection probabilities from these posteriors to understand the robustness of the results. It is now well established that parameters of neural models fall into two classes: those whose precise values are critical to the resulting predictions of a model, and those a model is not sensitive to (Panas et al., 2015; Ponce-Alvarez et al., 2020). And it is likely that striatal dynamics are indeed sensitive to variations in the probabilities and distances of connections (Humphries et al., 2010; Spreizer et al., 2017). Thus, we propose computational researchers change their usual practice of setting a single value for connection parameters, and instead sample from the posterior distribution on each run of their model – to this end, we give the complete form of all our posteriors for *p* in Table 1 and our confidence intervals for the decay parameter *β* in Table 3.

### Extending the microcircuit map

Constructing our map also revealed or reemphasised further research questions. First, because the available experimental data are drawn from across the striatum, the map is silent on anatomical issues such as whether connection probabilities differ between the patch and matrix compartments of the striatum or between different regions of the striatum. Second, we lack data on the connectivity of some types of striatal interneuron thought distinct to those examined here, including the calretinin-expressing interneurons and rare subtypes of 5HT3a-expressing interneurons (Tepper et al., 2018). Also omitted are the known connections from ACh interneurons to SPNs. As these synapses use muscarinic receptors, and so are metabotropic, they do not evoke postsynaptic currents detectable by the simultaneous stimulation and recording technique used in the studies relied upon here. However, new techniques such as induced overexpression of G-protein activated ion channels (Mamaligas and Ford, 2016) may allow the future quantification of connection rates. An advantage of our approach is that any new data on pairwise recording data in the striatum can build directly on our analyses, by either updating the posteriors we arrived at, or by estimating new posteriors for connections that lack data at present.

Third, we emphasise that this is a map of the local microcircuit, for the connections from a source neuron to other types of neurons in its neighbourhood. Our distance-dependent probability model assumes that connection probability falls monotonically with distance within the neighbourhood. However, sub-types of interneurons that send axons longer distances (Tepper et al., 2018) could violate this assumption, such as reports of PLTS interneurons with an axon that spans a distance of over 1mm making infrequent bouquets of terminals (Kawaguchi, 1993). More detailed knowledge of these long-distance connections would allow for a more complete map of striatal connectivity. Moreover, our approach requires a single parameter model of the distance-dependent probability, so assumes exponential decay…as p(connection) must decay over distance, this is fine; but more detailed….

Finally, this is a map of connectivity: full knowledge of the influence of one neuron type on another requires data on the strength of the different connections, which may in turn depend on where on the target neuron they fall (Oorschot et al., 2013; Du et al., 2017).

### Implications for theories of the striatum

Ever since the paper by Jaeger et al. (1994) which reported finding no functional lateral connections among SPNs, computational modelling of the striatum and wider basal ganglia moved away from earlier lateral inhibition models implementing a winner-takes-all strategy where these connections took centre stage (Groves, 1983) towards a predominantly feedforward view (Plenz, 2003; Tepper et al., 2004). According to these feedforward models (for example Mink (1996); Gurney et al. (2001b); Frank (2005); Leblois et al. (2006); Humphries et al. (2006)), the output of the striatum is entirely determined by the pattern of its cortical inputs modulated by the strength of the different cortico-striatal synapses.

Our Bayesian map of the striatal microcircuit provides further evidence that this feedforward model is limited because the striatum’s internal circuit is crucial in shaping its outputs. In the classic direct-indirect pathway model of the basal ganglia (Alexander and Crutcher, 1990), the D1 and D2 SPNs respectively form the populations originating the direct and indirect pathways to the basal ganglia output nuclei, D1 SPNs sending direct axonal projections, and the D2 SPNs projecting first to the globus pallidus, which relays to the output nuclei. We have shown that data on connections between D1 and D2 SPNs (Taverna et al., 2008; Planert et al., 2010) provides strong evidence for D2 SPNs making more connections to other SPNs than D1 SPNs. Some models and theories have interpreted these data as showing that the indirect pathway will dominate the direct pathway output (Bahuguna et al., 2015; Burke et al., 2017). However, that the same data provide no evidence of an asymmetry in the preferred targets of D1 or D2 SPNs further suggests there is not a selective inhibition of the D1 SPNs by D2 SPNs, but that the D2 SPNs are as equally likely to inhibit themselves as D1 SPNs. As such, the nature of the interaction of the direct and indirect pathways, and hence their response to cortical and thalamic inputs, remains unclear.

### Microcircuit mapping can be used anywhere in the brain

We showed a range of advantages that our Bayesian approach has over more traditional frequentist approaches. One is that it replaces a single point estimate surrounded by a flat confidence interval by a posterior distribution covering all the possible values, so that overlaps between connection rates become immediately apparent. Second, as argued in Dienes (2014), when differences between connection rates are non-significant, Bayesian methods allow us to distinguish between cases where the data is insufficient to draw a conclusion from cases where there really is no difference. For instance, the posteriors for connection rates in the study of Cepeda et al. (2013) strongly overlap (Figure 8), but because the posteriors are so broad, we know this is due to insufficient data rather than evidence of no difference. On the contrary, when we failed to find a difference in connection rates for different postsynaptic targets of the D1 and D2 SPNs, the posteriors are sufficiently narrow for us to confidently conclude that such a difference is either absent or quite small (Figure 2). Third, when making comparisons between probabilities of connection, we can use the full posteriors to compute an explicit probability that the difference is less than or greater than zero. A final advantage of Bayesian inference is the use of priors to incorporate past results, as we did for the connections between SPNs, or test our starting assumptions, as we did by showing our SPN connection results were robust to the choice of prior distribution.

Our approach can easily be applied to any brain regions where paired recording experiments have taken place, such as the recent study of Ellender et al. (2019) on how the embryonic origin of cortical neurons influences their connection probabilities. Indeed, obtaining the posterior distributions given *k* positive tests of connection and *n* – *k* negative tests requires a single line of MATLAB or Python thanks to built-in functions (Methods). Consequently, not only are these Bayesian methods easily applicable to intracellular recording data from any brain region, but also may be a rare case where it easier to be Bayesian than frequentist.

## ACKNOWLEDGEMENTS

This work was supported by the Medical Research Council [grant numbers MR/J008648/1, MR/P005659/1 and MR/S025944/1]. We thank Robert Schmidt and Benoît Girard for their helpful comments on a draft of this paper.

## Author Contributions

F.C. and M.D.H designed the analyses. F.C. analysed the data. F.C. and M.D.H wrote the manuscript.

## Notes

### Competing Interest Statement

The authors have declared no competing interest.

### Summary of Updates

Added a more detailed justification for teh choice of exponential decay as a model of distance-dependent connection rates (line 130-145)

